# Digital Chromosome Banding Reveals Distinct Spatiotemporal Dynamics and Sexual Dimorphism in Meiotic Silencing

**DOI:** 10.64898/2026.05.28.728562

**Authors:** Yiheng Peng, Shuangqi Wang, Royal Shrestha, Xing Tian, Evan Zakharchenko, So Maezawa, Carolline Ascencao, Marcus Smolka, Paula Cohen, Prabhakara Poothi Reddi, Satoshi H. Namekawa, Huanyu Qiao

## Abstract

In mammals, meiotic silencing of unsynapsed chromatin (MSUC) is initiated by the DNA damage response (DDR) pathway, as marked by γH2AX. During normal male meiosis, MSUC is restricted to the unsynapsed sex chromosomes, a process known as meiotic sex chromosome inactivation (MSCI). While the initiation of MSCI has been well studied, its full silencing dynamics and underlying structural mechanisms remain unclear. In contrast to MSCI, broader MSUC can occur on autosomes in response to synapsis failure, but its cell-to-cell variability obscures its quantification. To address these challenges, we introduce “digital-chromosome-banding”, a single-cell-based approach that allows quantitative analysis of MSCI and MSUC at chromosomal resolution. Using this approach, we identified two distinct silencing transitions during MSCI, occurring from zygonema to early pachynema and from early to mid-pachynema. The latter step coincides with mature sex body formation and involves a gel-like diffusion barrier to enforce transcriptional repression. Applying this approach to synapsis-defective mouse models (*Spo11^−/^*^−^, *Tardbp* cKO, and *Nelfb* cKO), we observed divergent MSUC patterns that correlate with the severity of asynapsis. Comparative analysis between sexes also uncovered notable sexual dimorphisms in meiotic silencing. Together, our data provide a quantitative framework to dissect the spatiotemporal dynamics and sexual differences of meiotic silencing.

## Introduction

Meiosis is a conserved and highly regulated process that generates haploid gametes -- sperm and oocytes in mammals. During meiotic prophase I, homologous chromosomes undergo pairing, synapsis, and recombination, events critical for accurate segregation and the promotion of genetic diversity^1^. Disruption in these events can impair germ cell development, triggering meiotic arrest and apoptosis^2,3^.

To safeguard meiotic progression, evolutionarily conserved checkpoint mechanisms monitor the fidelity of homologous synapsis and recombination. A key outcome of checkpoint activation is the transcriptional silencing of unsynapsed chromatin^4^. In males, this manifests as meiotic sex chromosome inactivation (MSCI), arising from asynapsis of the X and Y chromosomes^5^. Failure to initiate MSCI leads to meiotic arrest and spermatocyte elimination^6,7^. Mechanistically, MSCI initiation is orchestrated by DNA damage response (DDR) proteins such as Breast Cancer gene 1 (BRCA1) and Ataxia Telangiectasia and Rad3-related (ATR), which localize to the unsynapsed regions of the sex chromosomes in early pachynema to form the repressive sex body^8,9^. BRCA1 promotes ATR recruitment, which is activated by Topoisomerase 2 Binding Protein 1 (TOPBP1) and phosphorylates the histone variant H2AX (γH2AX) to initiate transcriptional silencing^10,11,12^. However, even though the initiation of MSCI has been well characterized, the mechanisms that sustain stable and complete silencing throughout prophase I remain poorly understood.

In mammals, MSCI represents a specialized form of the broader process known as meiotic silencing of unsynapsed chromatin (MSUC), which serves as a surveillance mechanism against chromosome asynapsis to maintain genome integrity^13,14,15^. MSUC is driven by DDR signaling, with γH2AX accumulation marking unsynapsed chromatin^13^. In the absence of proper synapsis, such as in *Spo11^−/−^* spermatocytes, MSUC occurs ectopically and form repressive nuclear domains known as pseudo-sex bodies^16^. Extensive studies have characterized MSUC using microscopy-based assays, such as Cot-1 RNA fluorescence in situ hybridization (FISH) and gene-specific RNA FISH^5,13,17^. These approaches provide high-resolution, locus-specific insights into transcriptional silencing. Transcriptomic approaches, including microarray, RNA-seq, and single-cell RNA-seq (scRNA-seq), have been effectively applied to quantify MSCI^18,19,20,21^. However, accurately quantifying MSUC at the genome-wide level remains challenging, particularly due to the stochastic and cell-to-cell heterogeneous nature of chromosomal asynapsis.

Here, we introduced a complementary and novel approach, termed “digital-chromosome-banding”, which enables quantitative, region-specific analysis of transcriptional repression at single-cell resolution, allowing systematical dissect of the spatial and temporal dynamics of both MSCI and MSUC. Applying this approach in wild-type spermatocytes, we found that MSCI proceeds in a stepwise manner throughout prophase I, suggesting that stable silencing is progressively established in coordination with chromatin reorganization. Further analysis of the *Topbp1^B5/B^*^5^ model^22^, which lacks proper sex-body maturation, confirmed that although MSCI initiation occurs, its progression to full silencing is impaired, implicating structural remodeling of the sex body as a key determinant of full MSCI. Consistent with this, we found that sex-body maturation involves the formation of a gel-like sex-chromosomal domain that restricts the diffusion of small molecules. Analysis of MSUC on three synapsis-defective mouse models (*Spo11^−/−^, Tardbp* cKO, and *Nelfb* cKO) displayed divergent silencing patterns. Together, these findings delineate distinct regulatory features of MSCI and MSUC, providing a quantitative foundation for understanding the spatial, temporal, and structural dynamics of meiotic silencing.

## Results

### Progressive silencing of MSCI was quantified by digital-chromosome-banding

To capture the temporal progression of MSCI silencing, we analyzed transcriptional dynamics in meiotic germ cells using scRNA-seq datasets because scRNA-seq can provide temporal stage information without cell sorting. Given that transcriptional silencing may occur unevenly across different chromosomal regions, such as the pseudoautosomal regions (PARs) versus the non-PAR, heterochromatic regions of the Y chromosome during pachynema, we developed a scRNA-seq-based approach to quantify transcriptional activity at a regional level. This method, termed digital-chromosome-banding, virtually resembles Giemsa chromosome banding (G-banding) and employs a bin-shifting strategy to systematically scan chromosomes (Figs. 1A,B). Specifically, we divided each chromosome into small (10Mb) bins and quantified transcription based on the cumulative expression of all genes within each bin (Fig. 1A). Transcription level of each chromosome across individual cells was visualized in a heatmap, with columns representing cells and rows corresponding to chromosomal bins (Fig. 1B).

**Fig. 1.**
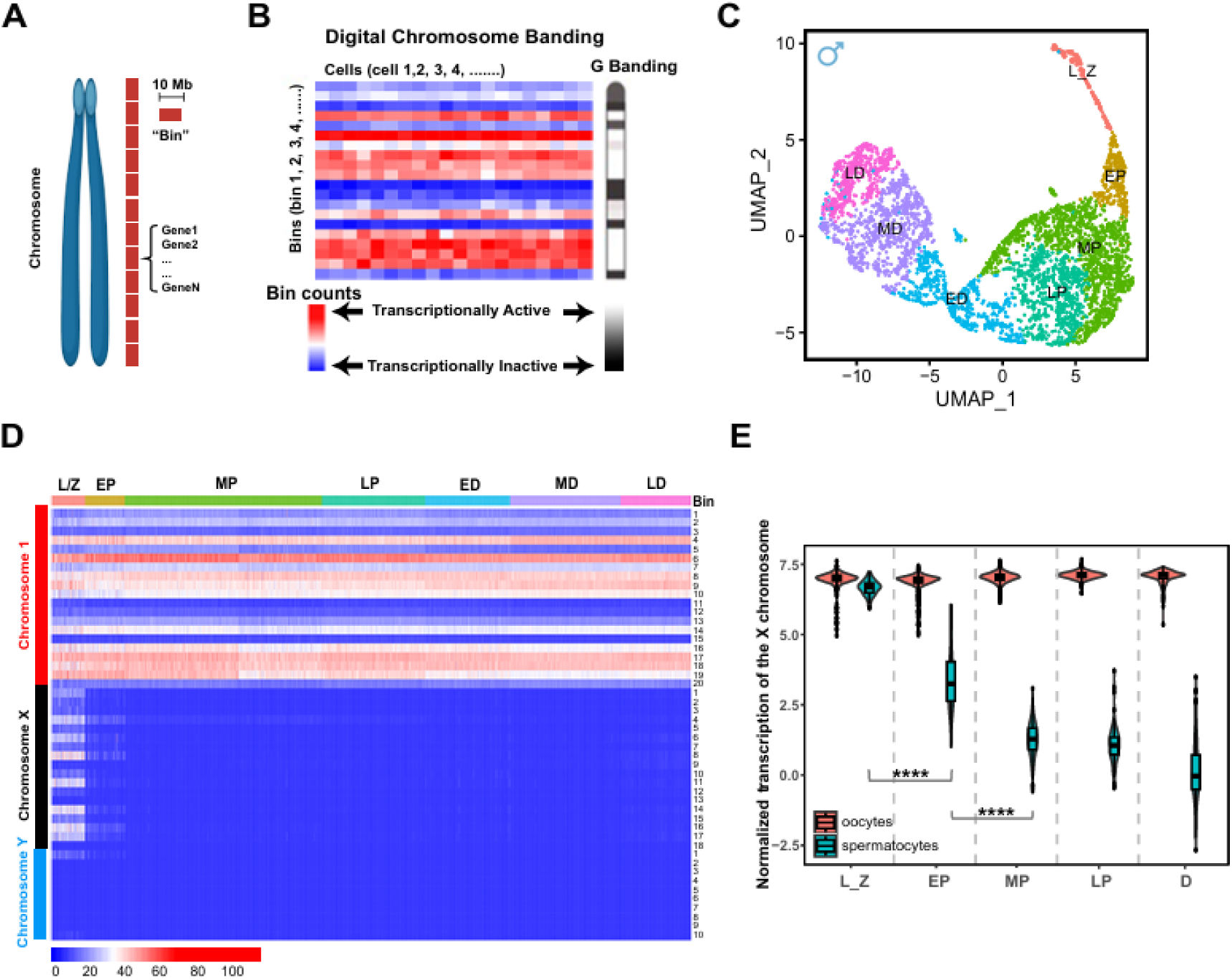
Digital-chromosome-banding illustrates transcriptional dynamics across meiotic prophase. **I.** (A) Schematic representation of the digital-chromosome-banding approach. Each chromosome is segmented into 10 Mb bins, with total transcripts per bin summed and log-transformed to represent their transcriptional activity. (B) Digital-chromosome-banding heatmap resembles cytological chromosome Giemsa (G) banding patterns. The heatmap displays bin transcriptional levels along chromosome 1 in wild-type pachytene spermatocytes. Rows represent bins, while columns correspond to individual cells. (C) UMAP analysis of spermatocyte clustering across meiotic prophase I. Cells are categorized into distinct stages: L_Z (leptotene_zygotene), EP (early pachytene), MP (mid pachytene), LP (late pachytene), ED (early diplotene), MD (mid diplotene), LD (late diplotene). (D) Digital-chromosome-banding heatmap of chromosomes X, Y, and 1 across prophase I. Generated from single-cell RNA-seq data (Fig. 1C), the heatmap reveals transcriptional silencing of X and Y chromosomes beginning at the early pachytene stage. (E) Violin and box plots showing relative transcriptional levels of X chromosome bins throughout meiotic prophase I in oocytes and spermatocytes. In spermatocytes, two distinct reductions in X chromosome transcription occur: from zygotene to early pachytene and from early pachytene to mid-pachytene. P-values were calculated using the Wilcoxon test; **** indicates p < 0.0001. Average expression levels of X chromosome: oocytes (119, 131, 137) and spermatocytes (15.0, 2.29, 2.15) in early, mid, and late pachytene, respectively, corresponding to 8-fold, 60-fold, and 64-fold reductions.

To investigate MSCI silencing patterns, we analyzed wild-type (WT) testicular scRNA-seq datasets from previous studies (Supplementary Table 1)^22^, and extracted 4010 spermatocytes in meiotic prophase I (Fig. 1C). Using Uniform Manifold Approximation and Projection (UMAP), we categorized these cells into leptotene/zygotene, pachytene, and diplotene stages. To monitor MSCI progress with a finer time resolution, we further subdivided pachytene and diplotene into early, mid, and late substages (Fig. 1C), defined by germ-cell-specific markers (Extended Data Fig. 1A). By applying digital-chromosome-banding to these meiotic cells, we examined transcriptional dynamics across prophase I. Notably, autosomes and sex chromosomes exhibited distinct temporal transcriptional dynamics (Figs. 1D,E). Autosomal transcription levels remained relatively stable throughout prophase I, whereas a substantial reduction in transcription was observed in the X chromosomes and PARs of the Y chromosomes after zygonema (Figs. 1D,E), consistent with previous reported MSCI timing^23,24^. Interestingly, additional transcriptional silencing of the X chromosomes was observed during the transition from early to mid-pachynema, suggesting that MSCI is a progressive process rather than a single transcriptional shutdown event.

To exclude the possibility that the observed stepwise reduction of X-linked transcripts was driven by RNA stability rather than transcriptional silencing, we performed two additional analyses focused on nascent transcription. First, we analyzed an independent single-nucleus RNA-seq (snRNA-seq) dataset from WT mouse testicular cells^19^. Ordering germ cells along spermatogenic progression revealed a clear reduction in X-linked transcript abundance in spermatocytes (Extended Data Fig. 2A,B), consistent with the onset of MSCI^25^. Second, to directly assess whether the stepwise reduction reflected ongoing transcriptional changes, we extracted and reanalyzed intronic reads from scRNA-seq datasets. Intronic reads are enriched for nascent pre-mRNA and therefore provide a proxy for newly synthesized transcripts. Total intronic RNA levels increased across spermatogenic stages, consistent with elevated global transcriptional activity during meiotic progression (Extended Data Fig. 2C) ^25^. In contrast, X-linked intronic RNA levels showed a marked and stepwise reduction, with decreases observed from zygonema to early pachynema and again from early to mid-pachynema (Extended Data Fig. 2D). These results indicate that the biphasic reduction of X-linked transcripts cannot be explained solely by differences in RNA stability, and instead support a model in which MSCI proceeds through progressive, stepwise transcriptional silencing during prophase I.

MSCI is a male-specific event driven by the incomplete synapsis of X and Y chromosomes. To quantify the silencing extent of MSCI, we compared X chromosome expression in spermatocytes versus oocytes where MSCI does not occur (Fig. 1E). We analyzed 5107 oocytes in meiotic prophase I as a control (Extended Data Fig. 3 and Supplementary Table 1)^26,27,28^. Our analysis revealed that, in early pachytene, the average transcriptional levels of spermatocytes were 8-fold lower than that of oocytes. By mid and late pachytene, transcriptional levels were reduced by 60-fold and 64-fold, respectively (Fig. 1E and Extended Data Fig. 4).

In summary, digital-chromosome-banding provides a powerful quantitative framework to measure the temporal and spatial dynamics of meiotic silencing. Our findings reveal that MSCI occurs in a stepwise manner as prophase I progresses.

### MSCI is a multi-step process that correlates with phases of sex-body maturation

MSCI is accompanied by the formation of the sex body, a specialized nuclear domain where the unsynapsed X and Y chromosomes are sequestered and undergo heterochromatinization which occurs at the mid pachytene stage^29^. In WT spermatocytes, the shape of sex chromosomes is initially rod-like at the early pachytene stage, but becomes a round shape when XY body formation occurs in the mid pachytene stage^9^. However, it remains unclear whether distinct stages of MSCI correspond to specific phases of sex-body maturation. To investigate this, we examined *Topbp1^B5/B^*^5^ spermatocytes^22^, which initiate MSCI but fail to achieve complete silencing, providing a model to assess whether disrupted MSCI is associated with sex-body-maturation defects.

*Topbp1^B5/B^*^5^ spermatocytes can reach pachytene stage, as evidenced by the detection of H1t signal accumulation^22^. Here, we utilized this scRNA-seq datasets and first assessed the extent of sex chromosome silencing in *Topbp1^B5/B^*^5^ spermatocytes by comparing them with WT spermatocytes. Using digital-chromosome-banding, we visualized 200 spermatocytes at each substages in both WT and *Topbp1^B5/B^*^5^ mutant (Figs. 2A,B). Our analysis revealed that *Topbp1^B5/B^*^5^ spermatocytes at the final pachytene substage (P3) exhibited significantly higher transcription than WT early pachytene spermatocytes, indicating that MSCI stalls at an intermediate stage, failing to undergo the additional silencing step observed in WT late pachytene cells (Fig. 2C).

**Fig. 2.**
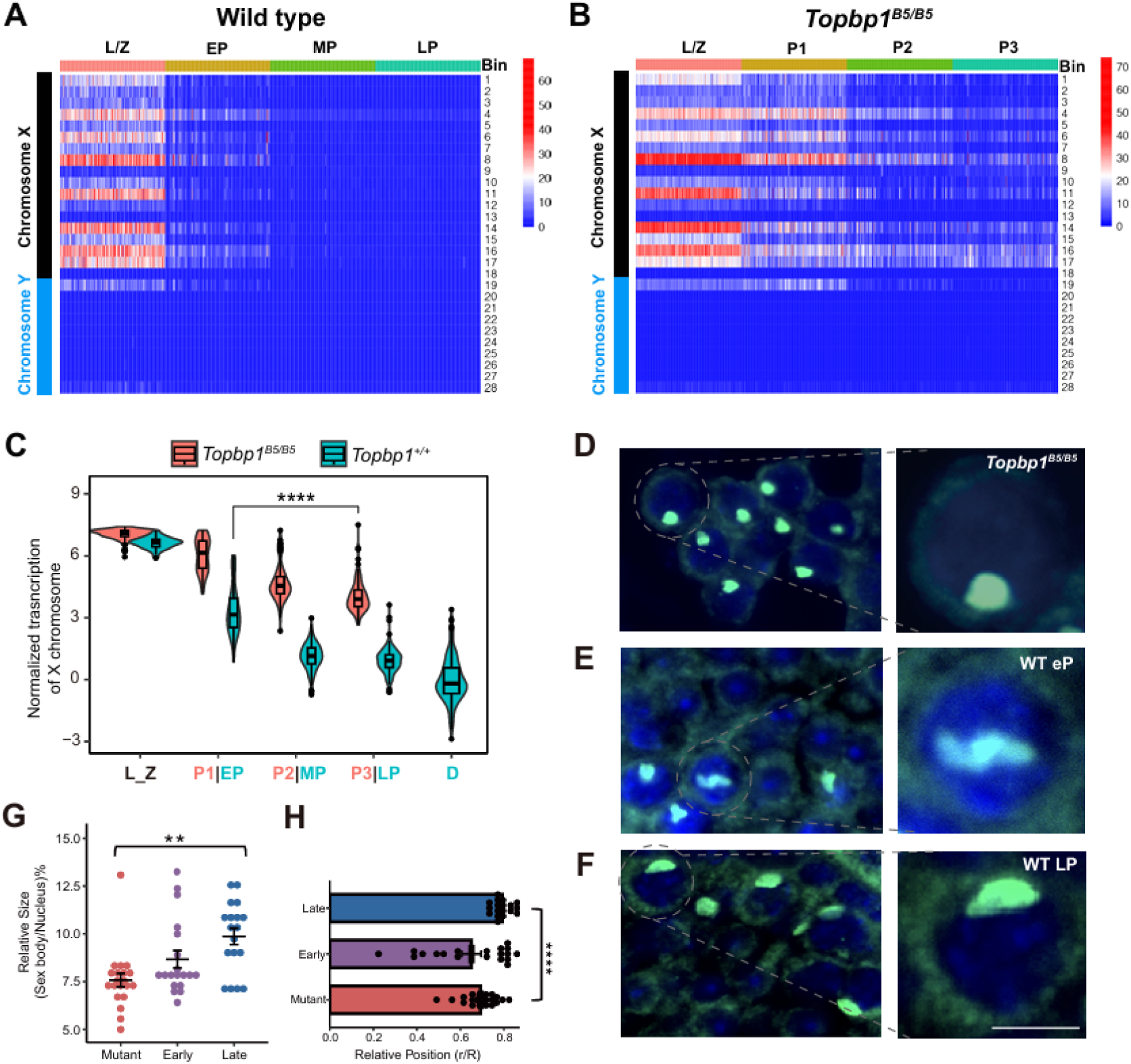
Incomplete X chromosome silencing and sex body organization in *Topbp1^B5/B5^* spermatocytes. (A-B) Digital-chromosome-banding heatmaps of wild-type (*Topbp1^+/+^*) spermatocytes (A) and *Topbp1^B5/B^*^5^ spermatocytes (B) during prophase I. L/Z: leptotene/zygotene; EP: early pachytene; MP: mid pachytene; LP: late pachytene; P1: pachytene stage 1; P2: pachytene stage 2; P3: pachytene stage 3. (C) Violin and box plot comparing the relative expression levels of bins along the X chromosome throughout meiotic prophase I in both *Topbp1^+/+^* and *Topbp1^B5/B^*^5^ spermatocytes. While MSCI initiates in *Topbp1^B5/B^*^5^ spermatocytes, it fails to reach to the first silencing step observed in wild-type MSCI during early pachynema. P-values were calculated using the Wilcoxon test. **** indicates p-value < 0.0001. (D-F) Immunostaining of histology testes sections from *Topbp1^B5/B^*^5^ spermatocytes (D), WT early pachytene (E), and WT late pachytene spermatocytes (F). Nuclei were stained with γH2AX (green) and counterstained with DAPI (blue). Scale bar = 25 μm. (G) Comparison of relative sex-body size in *Topbp1^B5/B^*^5^ spermatocytes, WT early and WT late pachytene spermatocytes. P-value were calculated using the Wilcoxon test. ** indicate p-value < 0.01. (H) Comparison of relative nuclear positioning of sex bodies in *Topbp1^B5/B^*^5^ spermatocytes, WT early and WT late pachytene spermatocytes. P-value were calculated using Wilcoxon test. **** indicate p-value < 0.0001.

In WT spermatocytes, as MSCI progresses, the sex body compacts and migrates toward the nuclear periphery (Extended Data Figs. 5 and 6). Previous study has demonstrated that *Topbp1^B5/B^*^5^ spermatocytes can form sex bodies^22^. However, since these spermatocytes failed to progress beyond the second major silencing step, we asked whether sex-body maturation was disrupted. To address this, we compared histological sections of testis tissue from *Topbp1^B5/B^*^5^ and WT pachytene spermatocytes, focusing on sex-body morphology and nuclear positioning (Figs. 2D-F). We quantified sex-body compaction by normalizing its area to nuclear size and found that *Topbp1^B5/B^*^5^ pachytene spermatocytes had significantly less condensed sex bodies than WT late pachytene spermatocytes (Fig. 2G). Additionally, *Topbp1^B5/B^*^5^ sex bodies remained centrally positioned within the nucleus, rather than migrating toward the periphery as seen in WT late pachytene spermatocytes (Fig. 2H). Similar observations were made in immunofluorescence staining of chromosome spreads (Extended Data Fig. 7). These results indicate although a sex body forms in *Topbp1^B5/B^*^5^, it was less condensed and remains more centrally positioned, suggesting incomplete isolation compared to that of WT late pachytene stage.

Together, the analysis of *Topbp1^B5/B^*^5^ spermatocytes indicates that sex-body maturation is functionally linked to the later stages of MSCI. The incomplete MSCI observed in *Topbp1^B5/B^*^5^ spermatocytes correlates with defective sex body condensation and positioning, highlighting the interdependence between MSCI progression and nuclear organization during meiosis.

### Mature sex body exhibits gel-like properties that repel small molecules

We have demonstrated that sex body maturation is a gradual process that parallels MSCI progression. Matured late-stage sex bodies are proposed to form a distinct transcriptionally inert compartment that excludes transcription factors^29,30^. However, the biophysical properties of matured sex bodies remain poorly understood. Sex bodies have mainly been studied via immunostaining on fixed chromosome spreads, which requires antibody penetration into the nuclei and involves harsh treatment that disrupts cellular integrity. To avoid this, we performed live-cell imaging of sex bodies using the newly established SCML2-mClover transgenic mouse line (Extended Data Fig. 8)^31^. The polycomb protein SCML2 concentrates in the matured sex body from mid pachytene stage^32^. Thus, the conjugated mClover fluorescent protein acts as an intracellular probe to visualize sex bodies in the late pachytene/diplotene stages in green.

To confirm whether mature sex bodies form condensates that exclude molecules, we analyzed the penetration kinetics of small DNA dyes in spermatocytes. We stained SCML2-mClover spermatocytes with Hoechst 33342 (0.1ug/ml), a small, live-cell permeable DNA dye. Images were captured at various time intervals (5 min, 15 min, 25 min) post-staining. Within 5 minutes, strong Hoechst staining was observed in the outer nuclear layers, while weak signals were seen in the mature sex-bodies (Fig. 3A). Weak signals were also detected in the inner nuclear regions, suggesting delayed Hoechst penetration into the nuclear center (Fig. 3A and Extended Data Fig. 9). After 15 minutes, Hoechst signal intensity increased in the sex body and inner nuclei, whereas the outer nuclei (excluding the sex body) remained unchanged. To quantify Hoechst penetration rate in the sex body, we calculated the relative Hoechst staining by dividing the fluorescence intensity of the sex-body region by the non-sex-body nuclear regions. In live spermatocytes, relative Hoechst staining increased largely after 15 and 25 minutes, suggesting delayed penetration into the sex-body regions. In contrast, Hoechst efficiently stained heterochromatin regions outside the sex body (Fig. 3B).

**Fig. 3.**
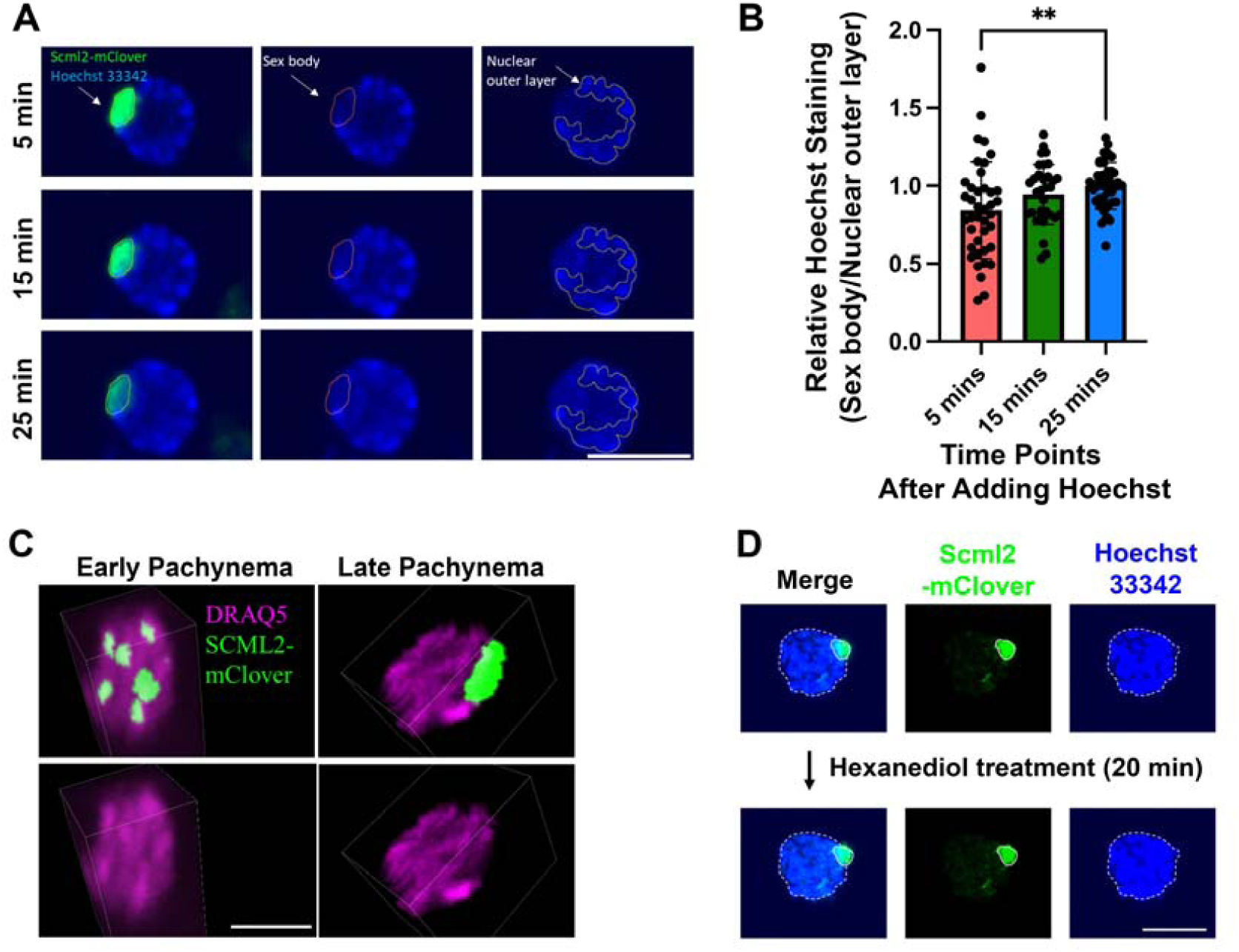
The mature sex body is a gel-like condensate that excludes small molecules. (A) Hoechst efficiently stained heterochromatin regions except sex bodies. A Hoechst 33342-stained SCML2-mClover-positive spermatocyte was imaged at different timepoints (5, 15, 25 minutes after Hoechst addition). The sex-body region and non-sex-body outer nuclear region were outlined with a red line and a yellow line, respectively. Scale bar = 25 μm. (B) Pairwise t-test for relative Hoechst staining at the timepoints in (B) (5, 15, 25 minutes after Hoechst addition, n = 40, 33, 42). Relative Hoechst staining is calculated by dividing the mean Hoechst intensity of the sex-body region by the mean intensity of the non-sex-body outer nuclear region. (C) 3D imaging after DRAQ5 treatment of SCML2-mClover-positive spermatocytes for 10 minutes. DRAQ5 and SCML2-mClover signals are shown for early-pachytene (left) and late-pachytene (right) spermatocytes. A video (Video 1) further demonstrates differences in DRAQ5 penetration within SCML2-mClover regions between early- and late-pachytene nuclei. (D) Effects of Hexanediol treatment on the sex body. Top row: Hoechst 33342 staining of SCML2-mClover-positive spermatocytes highlights the relative position of the sex body (SCML2-mClover, green) within the nucleus (Hoechst, blue). Bottom row: After imaging, the same spermatocyte was treated with 5% 1,6-Hexanediol for 20 min and a new image was taken using the same settings. The nucleys stained with Hoechst 33342 is outlined with a dashed white line and the sex body is highlighted with a solid white line. No disruption of sex-body shape was observed after Hexanediol treatment, suggesting that the sex body is not a permeable, liquid-like structure. Scale bar = 25 μm.

To further validate these findings, we repeated the experiment using another small, live-cell permeable DNA dye, DRAQ5. SCML2 is widely distributed in the nucleus before pachytene but concentrates in the sex body regions during pachytene stage. Therefore, the SCML2 signal pattern can be used to identify the maturation stage of the sex body^31^. A fully concentrated SCML2 signal represents a mature sex body, whereas a dispersed SCML2 signal indicates an immature sex body (Fig. 3C). Within 10 minutes of DRAQ5 treatment, mature sex body regions exhibited weaker DRAQ5 signals compared to other nuclear regions. In contrast, immature sex bodies exhibited DRAQ5 signal intensity comparable to other nuclear regions, indicating a difference in penetration velocity between mature and immature sex bodies (Fig. 3C and Video 1).

Depending on their density and physical state, mature sex bodies may adopt liquid-like, gel-like, or solid-like properties^33^. To assess their structural properties, we first tested whether they exhibit liquid-like structures, as suggested by Lin et al. (2024)^34^. 1,6-hexanediol is commonly used to disrupt cellular assemblies with liquid-like properties but is ineffective for non-LLPS (Liquid-Liquid Phase Separation) structures, such as nucleoli^35^. Our experiments showed that 1,6-hexanediol failed to disrupt the mClover fluorescence in sex bodies, indicating that sex bodies are not liquid-like structures (Fig. 3D). Furthermore, unlike amyloid, which form irreversible solid-like assemblies, sex bodies can disassemble. This observation suggests that they are more likely to be reversible gel-like structures rather than irreversible solid-like structures. Thioflavin T (ThT) is a dye that stains protein aggregates. If sex bodies exhibited solid-like properties, ThT fluorescence would be expected in sex-body regions. However, ThT stained the pachytene nuclei but excluded sex bodies, confirming that sex bodies are not solid-like structure (Extended Data Fig. 10). Our results suggest, albeit indirectly, that mature pachytene/diplotene sex bodies assemble into a compact, more likely gel-like structure that impedes rapid penetration of small molecules like Hoechst and DRAQ5.

### Detection of MSUC in *Spo11^−/−^* spermatocytes

MSUC is a more general transcriptional silencing mechanism targeting unsynapsed chromatin. To study this broader response, we used *Spo11*^−/−^ spermatocytes, where the absence of meiotic DSBs leads to genome-wide asynapsis and triggers MSUC^36^. *Spo11*^−/−^ spermatocytes form the pseudo-sex body resembling the inactive sex body seen in MSCI. However, unlike the X/Y-restricted sex body, the pseudo-sex body in *Spo11*^−/−^ spermatocytes can occur randomly. To quantify MSUC associated with asynapsis, we generated scRNA-seq libraries from *Spo11*^−/−^spermatocytes. Among 4283 testicular cells, 274 spermatocytes were identified based on high expression of germ-cell specific markers such as *Dazl*, *Sycp1*, *Sycp3*, *Smc1b* (Fig. 4A and Extended Data Fig. 11). Interestingly, we identified a subset of germ cells (n=75) exhibiting spermatid-like transcriptional signatures, characterized by the high expression of *Prm1* and *Prm2* (protamine proteins essential for sperm chromatin condensation)^37^ as well as *Tnp1* and *Tnp2* (expressed in round and elongating spermatids and involved in histone-to-protamine replacement)^38^ (Figs. 4B,C and Extended Data Fig. 12A). Since *Spo11*^−/−^ spermatocytes fail to progress beyond pachynema^39^, the presence of these cells is unexpected. We refer to them as mutant/“spermatid-like” (SPT-like) cells, as they exhibit transcriptomic features beyond early meiotic stages and thus resemble post-meiotic spermatids.

**Fig. 4.**
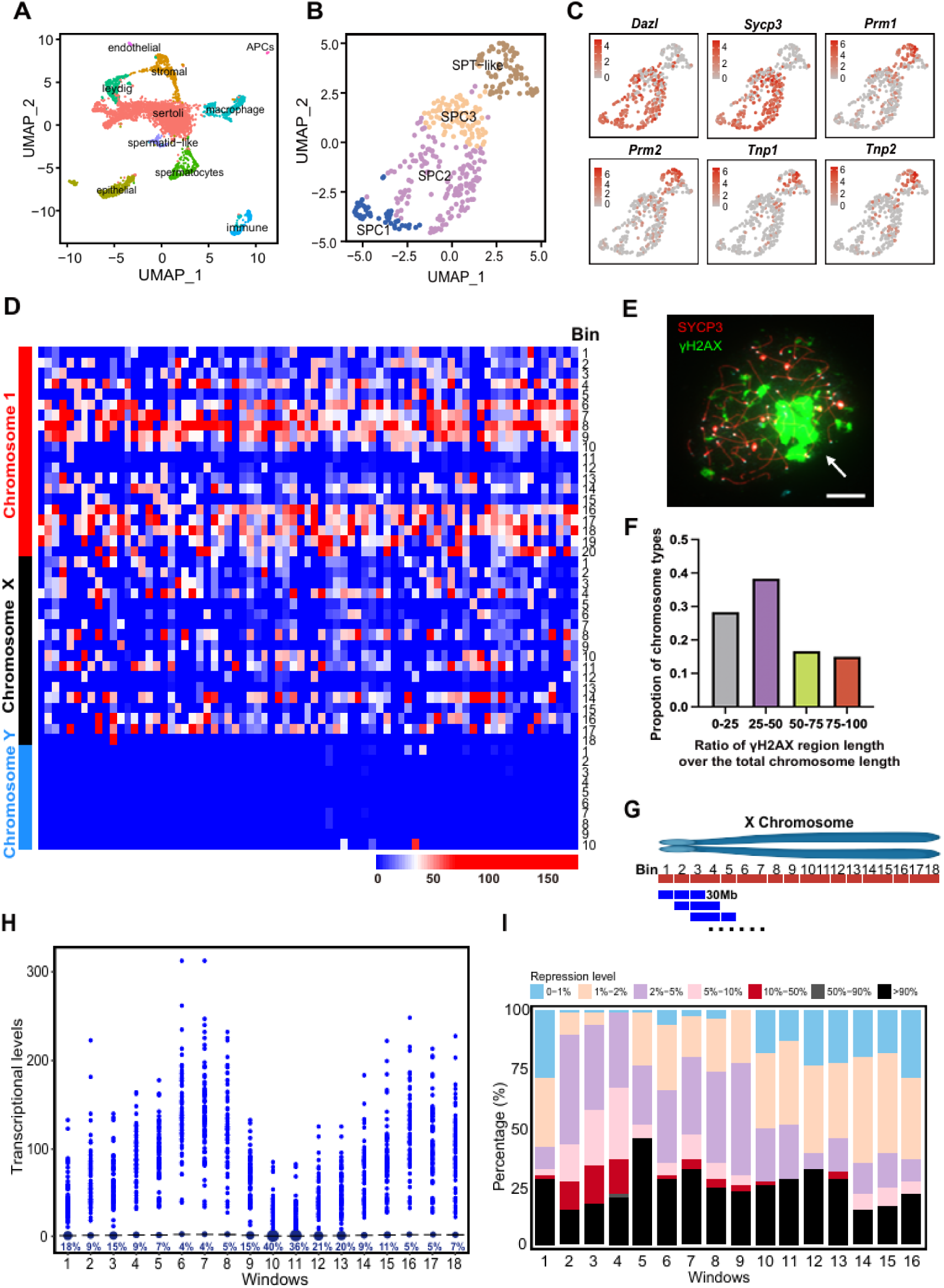
MSUC is detected in *Spo11*^−/−^ spermatocytes using digital-chromosome-banding. (A) UMAP plot of *Spo11*^−/−^ testicular cells. The identified cell types include Sertoli cells, stromal cells, epithelial cells, spermatocytes, Leydig cells, macrophages, immune cells, spermatid-like cells, endothelial cells, and antigen presenting cells (APCs). (B) UMAP plot of *Spo11*^−/−^ germline cells highlighting different cell stages. SPC: spermatocytes; SPT-like: spermatid-like cells. (C) UMAP plot of *Spo11*^−/−^ germline cells displaying expression of key marker genes: germ-cell-specific (*Dazl*), prophase I-specific (*Sycp3*), and spermatid-specific (*Tnp1/2*, *Prm1/2*). (D) Heatmap of transcriptional activity across 10 MB bins along chromosomes 1, X, and Y in *Spo11*^−/−^ SPT-like cells. Each column represents a cell, and each row represents a 10 Mb bin. (E) A representative immunofluorescent image of *Spo11*^−/−^ spermatocyte. γH2AX is in green and SYCP3 is in red. A white arrow marks a pseudo-sex-body region. Scale bar = 10 µm. (F) Bar plot quantifying chromosome involvement in pseudo-sex-body formation in *Spo11*^−/−^spermatocytes. Chromosomes are categorized based on the proportion of their length engaged in pseudo-sex-body formation: “0-25%”, “25-50%”, “50-70%”, and “75-100%” (X axis). Note: chromosomes without pseudo-sex-body involvement were excluded from this analysis. (G) Digital-chromosome-banding diagram, using a 30 Mb window with a 10 Mb shift. (H) Graph illustrating transcriptional levels of each 3-bin window along chromosome 1 in *Spo11^− /-^* SPT-like cells. Each dot represents an individual cell. The MSCI full silencing level (100% repression) is defined by the average transcriptional level of WT late pachytene spermatocytes (the stage when full silencing occurs). Black lines indicate MSCI levels for each window. The percentage of cells showing MSCI-like severe silencing is labeled in dark blue beneath the black MSCI threshold line for each window. Notably, MSCI-like severe silencing is observed across all 18 windows. (I) Stacked bar plot showing the percentage of transcriptional repression levels in *Spo11*^−/−^ SPT-like cells, measured across 16 consecutive 30 MB windows along the X chromosome. Each bar represents a 3-bin (30 MB) window, with colors indicating different levels of transcriptional repression. The black bars indicate high-level meiotic silencing is present in *Spo11*^−/−^ SPT-like cells.

To confirm MSUC occurrence in *Spo11*^−/−^ SPT-like cells, we applied digital-chromosome-banding, enabling transcriptional analysis at both regional and single-cell levels (Extended Data Fig. 13). Due to the random nature of pseudo-sex body, we expected to observe a “mosaic” expression pattern reflecting variability in transcriptional silencing across cells within the same chromosomal bin. As anticipated, the results revealed a heterogeneous transcriptional landscape, with “mosaic” silencing observed in both X chromosomes and autosomes, suggesting variable transcriptional repression among *Spo11*^−/−^ SPT-like cells (Fig. 4D).

To characterize MSUC, we first estimated the chromosome regions affected by MSUC. γH2AX immunostaining of *Spo11*^−/−^ spermatocytes revealed that ∼72% of chromosomes involved in pseudo-sex-body formation had γH2AX signals covering at least 25% of their length (Figs. 4E,F). Chromosomes with less than 25% involvement in pseudo-sex-body formation may not undergo stable silencing, considering the rapid chromosome movement during prophase I. Since the average mouse chromosome size is 124 Mb^40^, we estimated that ∼30Mb (124Mb × 25%) represents the minimal chromosomal region affected by MSUC. Based on this, we applied digital-chromosome-banding with a 30 Mb window and 10 Mb shift to systematically analyze transcriptional silencing in *Spo11*^−/−^ SPT-like cells (Fig. 4G). Analysis of transcriptional repression across 16 consecutive 30 Mb windows along the X chromosome confirmed significant meiotic silencing (Fig. 4I). Repression levels varied by region, with 17% - 45% of cells showing severe repression (> 90% silencing) per window. Similar analysis of chromosome 1, representing autosomes, showed that MSUC is not uniform but regionally variable, with 4%-40% cells reaching MSCI-like repression levels (Fig. 4H). Together, these results demonstrate the extent of pseudo-sex body-mediated transcriptional silencing in *Spo11*^−/−^ spermatocytes, highlighting the variability and regional specificity.

### MSUC patterns in *Spo11*^−/−^ spermatocytes

To further characterize MSUC in *Spo11^−/−^* spermatocytes, we analyzed transcriptionally active regions, as MSUC primarily affects these areas. Specifically, we examined active bins (represented by bin 4, bin 8, and bin 16) to assess transcriptional changes (Figs. 5A-D). In WT late pachytene spermatocytes, transcription within active bins of the X chromosome is uniformly silenced, reflecting the expected progression of MSCI. In contrast, autosomal transcription remains consistently high, maintaining normal transcriptional activity (Figs. 5A,B). However, *Spo11^−/−^* SPT-like cells exhibit a distinct bimodal transcriptional distribution within these active bins. This suggests that different cell populations exhibit variable transcriptional repression. Due to the stochastic nature of pseudo-sex-body formation, different cells may silence different regions randomly, leading to cell-to-cell variability in transcriptional repression. Consequently, an active bin may be fully silenced in some cells while remaining transcriptionally active in others.

**Fig. 5.**
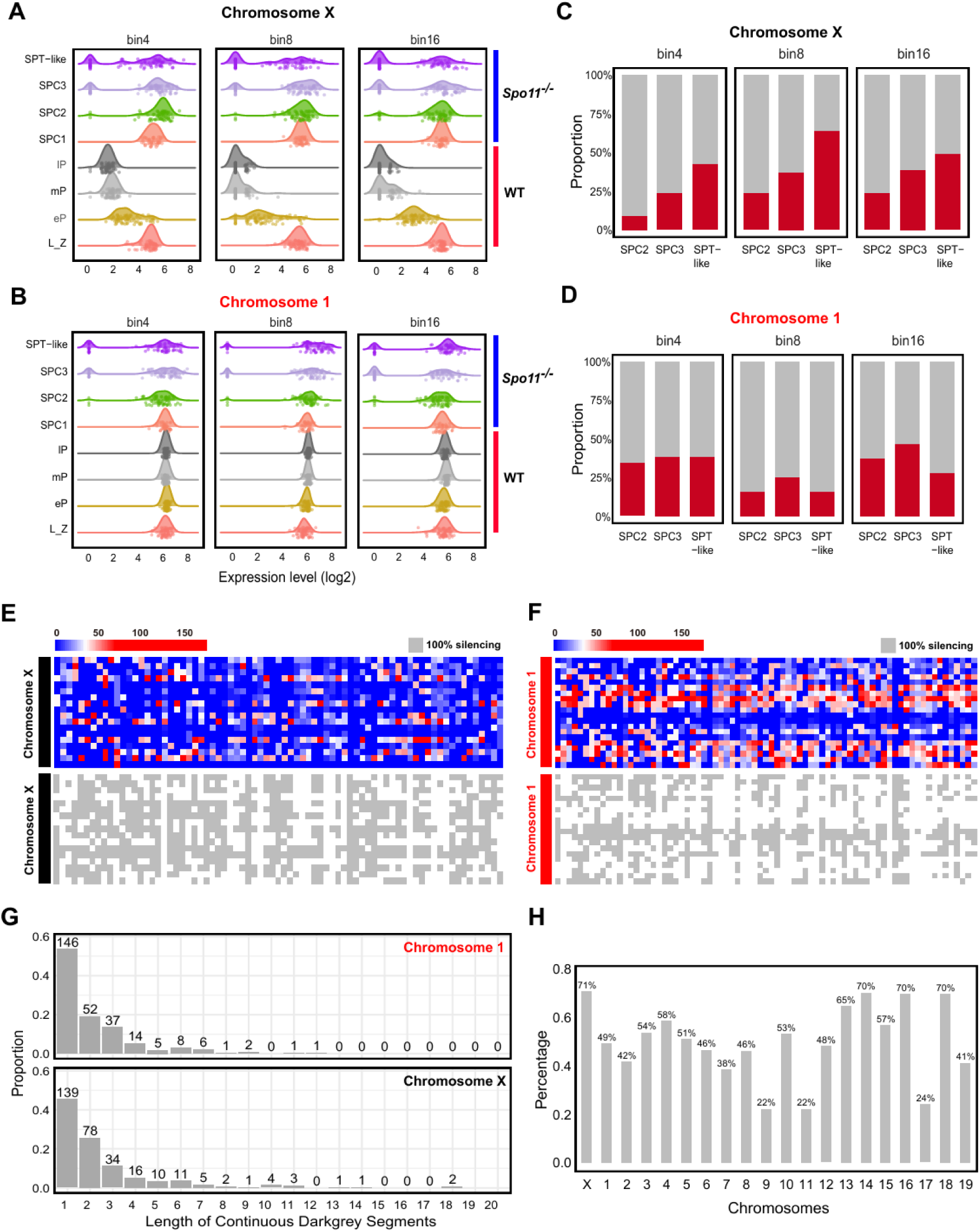
MSUC patterns across different stages and chromosomes. (A-B) Density plots illustrating transcriptional distribution of wild-type (WT) and *Spo11^−/−^*spermatocytes in active bins (represented by bin 4, bin 8, and bin 16) on chromosome X (A) and chromosome 1 (B) across different stages (rows). The x-axis represents log-transformed transcriptional level. Each dot indicates an individual cell. (C-D) Stacked bar plots showing the proportion of Spo11^−/−^ spermatocytes exhibiting MSCI-like silencing in active bins (bin 4, bin 8, and bin 16) on chromosome X (C) and chromosome 1 (D) across different stages. MSCI-like silencing is defined as transcription levels falling below the 5th percentile threshold of SPC1 spermatocytes; silenced portions are marked in red. (E-F) Heatmaps (top panels) showing the transcriptional activity for each 10 Mb bin along chromosome X (E) and chromosome 1 (F) among Spo11^−/−^ SPT-like cells. Grey-white heatmaps (bottom panels) display full silencing (grey) and active region (white). Columns indicate cells, and rows correspond to 10 Mb bins. (G) Bar plot showing the proportion of the continuous silenced regions of varying lengths along chromosomes X and 1, as seen in panel E-F. Proportions are labeled above each bar. (H) Bar plot indicating the proportion of Spo11^−/−^ SPT-like cells exhibiting MSUC across different chromosomes.

**Fig. 6.**
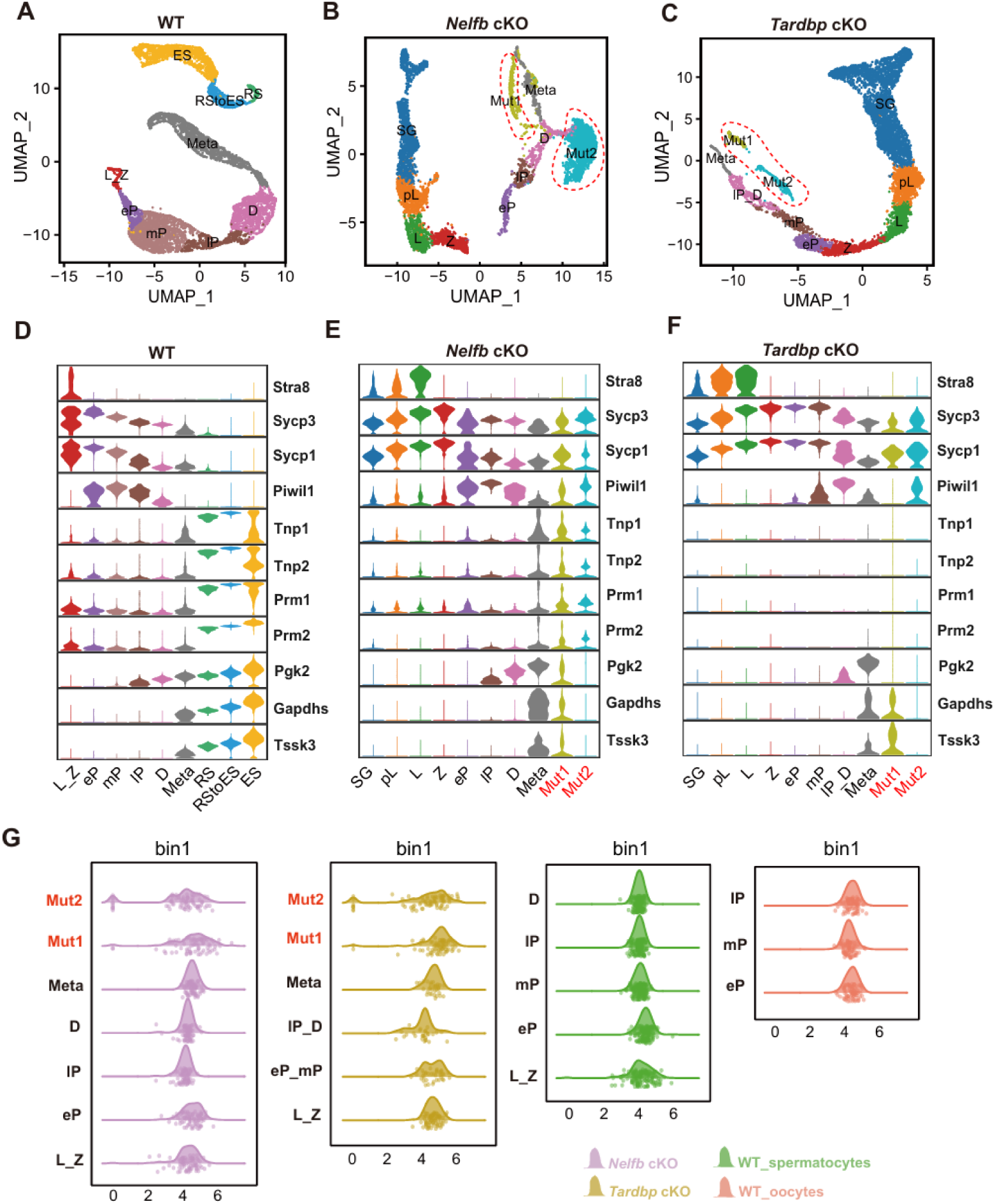
MSUC is detected in *Nelfb* cKO and *Tardbp* cKO germline cells via digital-chromosome-banding. (A-C) UMAP analysis of germline cells across different genotypes and developmental stages. (A) wild type, (B) *Nelfb* cKO, (C) *Tardbp* cKO. Spermatogenesis stages are denoted as SG: spermatogonia; pL: pre leptotene; L: leptotene; Z: zygotene; eP: early pachytene; mP: middle pachytene; lP: late pachytene; D: diplotene; Meta: metaphase; RS: round spermatid; ES: elongated spermatid. In both *Nelfb* and *Tardbp* cKO data sets, two transcriptionally distinct clusters deviating from the main developmental trajectory are marked with red dashed circles and designated as mutant populations. (D-F) Expression patterns of stage-specific marker genes across germline cell types in (D) wild-type, (E) *Nelfb* cKO, and (F) *Tardbp* cKO. (G) Density plots showing transcriptional distribution of cells across a representative bin (bin 1 on chromosome 6) of autosomes. The X axis represents log-transformed transcriptional levels. Each dot indicates an individual cell. Cell types are color-coded as follows: *Tardbp* cKO spermatocytes (light purple), *Nelfb* cKO spermatocytes (light yellow), wild-type spermatocytes (light green), and wild-type oocytes (light red).

To quantify this variation, we measured the percentage of *Spo11^−/−^*spermatocytes exhibiting full (MSCI-like) silencing in selected active bins of their X chromosome and autosomes. In the three examined X chromosome bins, we observed 43%, 64%, and 49% of cells showing MSCI-like silencing in SPT-like cells, whereas chromosome 1 (representing autosomes) displayed a lower extent of silencing, with 39%, 16%, and 28% of SPT-like cells exhibiting MSCI-like repression (Figs. 5C,D). Next, we measured silencing length by visualizing the transcriptional states using a grey-white heatmap, where grey represents full silencing (Figs. 5E,F). Among 75 *Spo11^−/−^* SPT-like cells, we identified 88 continuous grey segments (≥3 consecutive bins: 30 Mb) on the X chromosome, while 75 continuous grey segments were detected on chromosome 1 (Fig. 5G). To further quantify MSUC occurrence, we defined cells with at least three consecutive silenced bins (≥30 Mb, as estimated from immunostained spread analysis in Figs. 4E,F) as MSUC-positive. This analysis revealed that chromosomes X, 14, 16, and 18 were the most frequently involved in MSUC, with 70–71% of SPT-like cells exhibiting silencing in these chromosomes. Chromosomes 9, 11, and 17 were the least affected, with only 22–24% of cells showing MSUC (Fig. 5H).

In summary, our digital-chromosome-banding approach computationally defines MSUC transcriptional silencing patterns in *Spo11^−/−^*spermatocytes, characterizing the extent, segment length, and frequency of silencing across chromosomes. Our findings reveal a non-uniform, cell-specific pattern, supporting a stochastic model of pseudo-sex-body formation.

### Detection of MSUC in *Tardbp* and *Nelfb* cKO spermatocytes

To further explore MSUC in other asynapsis-prone models, we analyzed scRNA-seq datasets from *Tardbp* (gene encoding TDP-43) *and Nelfb* cKO spermatocytes, both of which are known to exhibit synapsis defects^41^.

Based on germ cell-specific markers (Extended Data Figs. 1C and 12B, C), we identified spermatocytes at different meiotic stages (Figs. 6A-C). In WT controls, cells followed a relatively continuous UMAP developmental trajectory, reflecting normal meiotic progression (Fig. 6A). In contrast, both *Tardbp* cKO and *Nelfb* cKO spermatocytes displayed two distinct clusters deviating from this trajectory, indicating transcriptional profiles that did not correspond to any defined meiotic stage. We refer to these populations as “Mut1” and “Mut2” (Figs. 6B, C). Interestingly, in *Nelfb* cKO spermatocytes, cells within these divergent mutant clusters expressed early meiotic genes like *Sycp1* and *Sycp3*, core components of the synaptonemal complex (SC) (Fig. 6B). Additionally, these cells also expressed the pachytene-stage-specific marker *Piwil1*, as well as post-meiotic genes such as *Prm1*, *Prm2*, *Tnp1*, and *Tnp2*, forming a transcriptional signature resembling spermatid-like cells—an observation also noted in *Spo11*^D*/*D^ mutants (Figs. 6D, E). Given that *Nelfb* cKO spermatocytes are known to arrest at the pachytene stage^41^, we hypothesize that the Mut1 and Mut2 clusters primarily represent pachytene-like mutant cells. Similarly, the undefined clusters in *Tardbp* cKO spermatocytes also expressed *Piwil1*, suggesting that these cells may also represent arrested pachytene-like cells resulting from asynapsis (Fig. 6F).

To assess whether MSUC occurs in *Nelfb* cKO and *Tardbp* cKO pachytene-like mutant spermatocytes, we performed digital-chromosome-banding. As expected, our analysis found that active bins (represented by bin 1 on chromosome 6) of autosomes exhibited a distinct bimodal transcriptional distribution (Fig. 6G), consistent with the observation in *Spo11^−/−^* SPT-like spermatocytes (Fig. 5A). This bimodal pattern likely reflects the random silencing nature of MSUC.

### MSUC-mediated silencing correlates with the degree of asynapsis

To assess the relationship between asynapsis and MSUC-mediated silencing, we first measured asynapsis levels via SYCP3 immunostaining in three synapsis-defective mutant models: *Tardbp* cKO, *Nelfb* cKO, and *Spo11^−/−^*. Compared to WT late pachytene spermatocytes, both *Tardbp* and *Nelfb* mutants showed moderate asynapsis, while *Spo11^−/−^* mutants exhibited a near-complete failure of chromosome synapsis (Fig. 7A). Quantification confirmed that *Tardbp* and *Nelfb* mutants had significantly fewer synapsed chromosomes than WT, with *Spo11^−/−^* displaying the most severe synaptic defects (Fig. 7B).

**Fig. 7.**
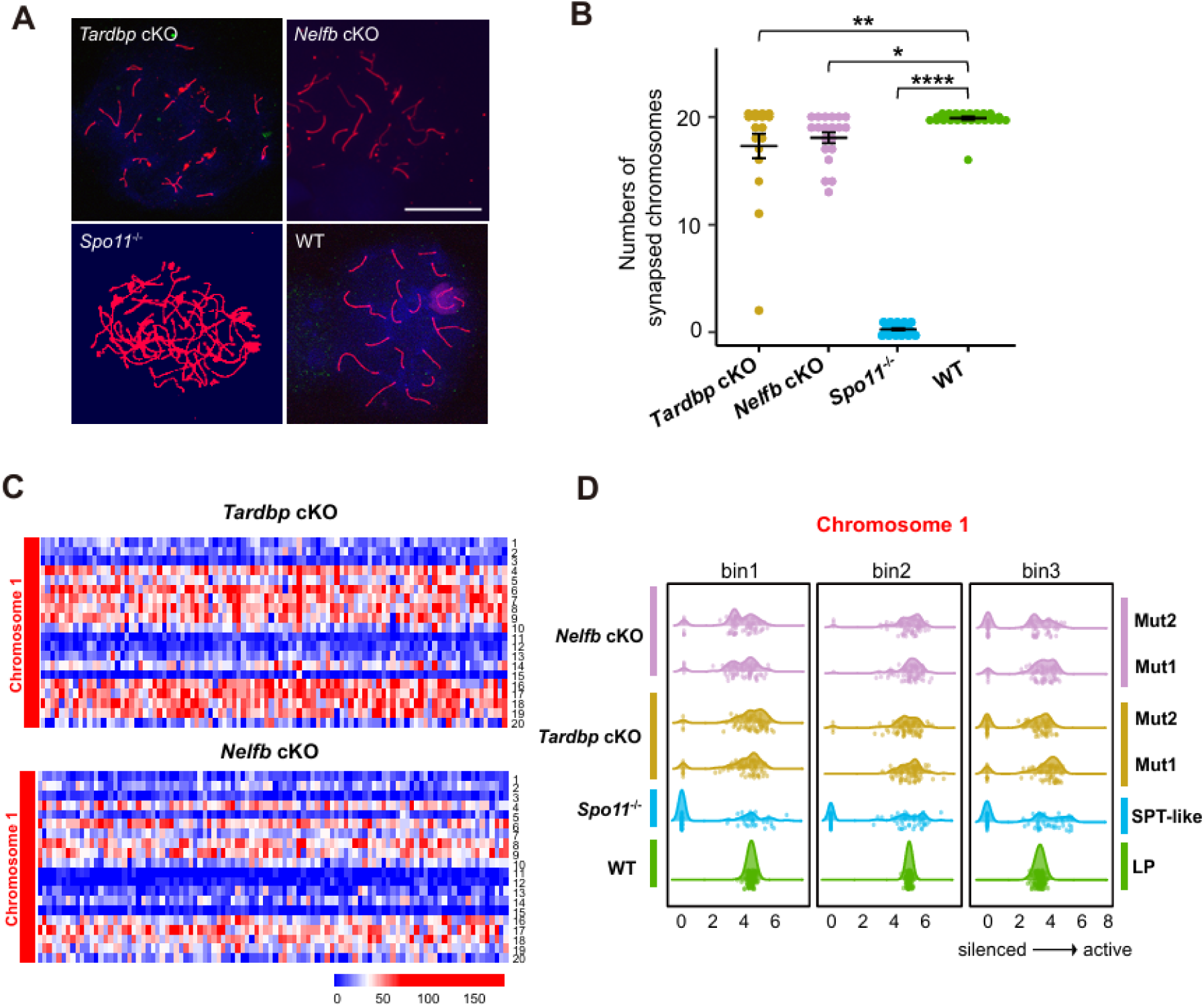
MSUC-mediated silencing is positively correlated with asynapsis levels. (A) Immunostaining of chromosome spreads from *Tardbp*, *Nelfb* cKO, and *Spo11^−/−^* mutant spermatocytes, as well as WT late pachytene spermatocytes. Nuclei were stained with SYCP3 (red) to visualize the axial elements (AEs) of the chromosomes. Scale bar = 20 μm. (B) Quantification of the numbers of synapsed chromosomes in mutant and WT spermatocytes. *, **, **** indicates p-value < 0.05, 0.01, 0.0001, respectively. (C) Heatmap of transcriptional activity across 10 Mb bins along chromosomes 1 in (up) *Tardbp* cKO, and (down) *Nelfb* cKO mutant spermatocytes. Each column represents a cell, and each row represents a 10 Mb bin. (D) Density plots showing transcriptional distribution of cells across bins (represented by bin 1-3) of chromosome 1 in *Nelfb*, *Tardbp*, *Spo11^−/−^* mutant spermatocytes, and WT late pachytene spermatocytes. The x-axis represents log-transformed transcriptional levels, where values from 0 to 8 indicate increasing transcriptional activity (from silenced to active). Each dot represents an individual cell.

We next examined transcriptional levels to evaluate the extent of MSUC. In the partially asynapsed *Tardbp* cKO and *Nelfb* cKO spermatocytes, transcriptional silencing diverged from both WT and *Spo11 ^/^ .* WT late pachytene spermatocytes showed full silencing (Fig. 1D), whereas *Spo11 ^/^* spermatocytes displayed mosaic repression across autosomes (Fig. 4D). In contrast, *Tardbp* and *Nelfb* cKO mutant spermatocytes exhibited noticeably slight transcriptional repression on autosomes represented by chromosome 1 (Fig. 7C). To quantify these differences, we generated density plots of transcriptional activity across representative 10 Mb bins of the chromosome 1 (bins 1-3). Although *Spo11^−/−^* spermatocytes showed a noticeable silencing peak on chromosome 1, *Tardbp* and *Nelfb* mutants exhibited only weak or even absent silencing signals in the same regions (Fig. 7D), suggesting a positive correlation between the severity of asynapsis and the extent of transcriptional silencing. Specifically, *Spo11^−/−^* shows the most extensive synapsis failure and also exhibits the strongest MSUC on autosomes. *Tardbp* and *Nelfb* mutants, with moderate synapsis defects, trigger a noticeably weaker, more limited MSUC. Together, these results demonstrate that the extent of MSUC-mediated silencing scales with the degree of asynapsis.

### High-level meiotic silencing is absent in pachytene oocytes

Similar to *Tardbp* cKO and *Nelfb* cKO male mutants that exhibit mild asynapsis, approximately 20% of wild-type oocytes in late prophase I also display partial asynapsis and form pseudo sex bodies marked by γH2AX immunostaining (Fig. 8A)^42^. Whether these asyanapsis regions in oocytes undergo MSUC-mediated gene silencing has been a subject of long-standing debate. It has been proposed that MSUC-mediated suppression of essential genes could trigger apoptosis, potentially contributing to the considerable loss of oocytes around birth^13^. However, other studies demonstrated that MSUC only plays a limited role in oocyte quality control, primarily eliminating oocytes with fewer than three pairs of asynapsed homologous chromosomes. In contrast, when asynapsis is extensive, reduced BRCA1 accumulation on each involved chromosome compromises MSUC initiation^43^.

**Fig. 8.**
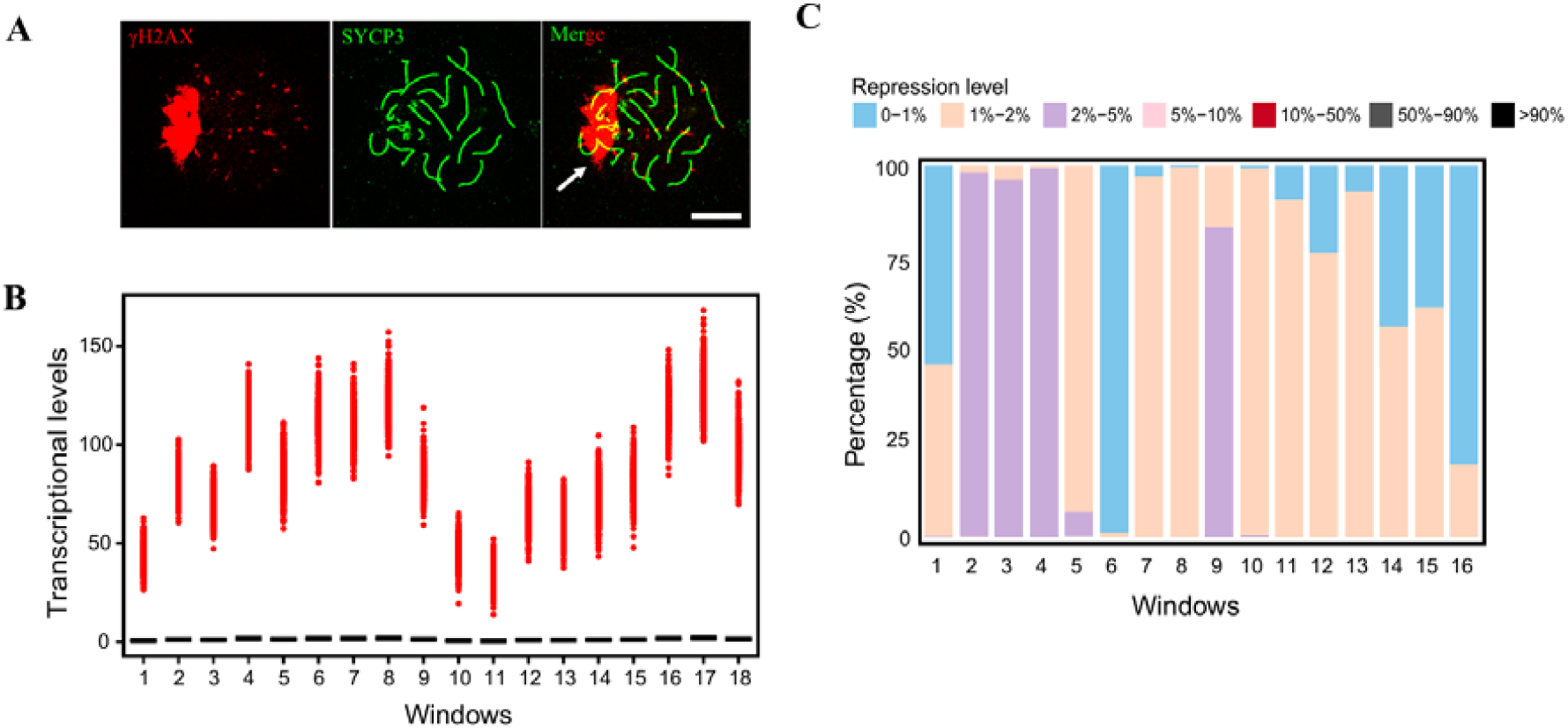
MSUC does not occur in oocytes despite asynapsis. (A) A representative immunofluorescent image of a late-pachytene oocyte showing γH2AX (red) and SYCP3 (green). A white arrow marks a pseudo-sex-body region. Scale bar = 10 µm. (B) Graph illustrating transcriptional levels of each 3-bin window along chromosome 1 in late pachytene oocytes. Each dot represents an individual cell. The MSCI full silencing level (100% repression) is defined by the average transcriptional level of WT late pachytene spermatocytes, when full silencing occurs. Black lines indicate MSCI levels for each window. Notably, MSCI-like severe silencing is absent across all 18 windows. (C) Stacked bar plot showing the percentage of transcriptional repression levels in late pachytene oocytes, measured across 16 consecutive 30 MB windows along the X chromosome. Each bar represents a 3-bin (30 MB) window, with colors indicating different levels of transcriptional repression. The absence of black segments indicates that high-level meiotic silencing is not present in late pachytene oocytes.

To assess the presence of MSUC-related silencing in wild-type oocytes, we scanned oocyte late-pachytene X chromosome with 30 MB (three bins) windows to detect MSCI-like silencing. Surprisingly, our analysis across 456 late pachytene oocytes failed to find any severe MSCI-like silencing on autosomes (represented by chromosome 1, Fig. 8B), and the X chromosome (Fig. 8C). Most late pachytene oocytes exhibited minimal silencing (< 2%), with only a few showings slightly a higher silencing level (2%-5%, e.g., the window 2-4 from chromosome X) (Fig. 8C).

Such low-level silencing patterns cast doubt on their association with MSUC. Because MSUC is a pachytene-specific event, we further examined whether such low-level silencing is stage-specific. Thus, we analyzed silencing patterns across different prophase I substages, including leptotene, zygotene, early pachytene, and mid pachytene stages (Extended Data Fig. 14A-D). Surprisingly, all stages exhibited similar low-level silencing patterns, suggesting that these effects are unrelated to MSUC. Together, our data strongly suggests the absence of MSUC in oocytes – an unexpected result that contrasts sharply with spermatocytes, where even mild asynapsis induces robust MSUC (Extended Data Fig. 14E-F).

## Discussion

In this study, we established digital-chromosome-banding, to directly and computationally detect transcriptional silencing in individual germline cells. This method overcomes the limitations of traditional approaches such as RNA FISH^13^, and bulk RNA-seq to assess MSUC^44^. In our study, by taking advantage of single-cell resolution and quantifying transcription across defined chromosomal subregions, our method can effectively resolve the variable and sporadic features of MSUC.

A key insight from our analysis is that MSCI silencing is not an instantaneous event, but instead follows a temporally structured, stepwise progression. We identified two major transcriptional shutdown phases: (1) the first during the zygotene to early pachytene transition, and (2) the second from early to mid-pachytene stage. This two-step silencing pattern temporally aligns with proposed models of MSCI regulation, in which DDR factors are initially recruited to sex chromosomes at early pachytene and later sequestered within the XY body from mid-pachytene onward^9^. The correlation between transcriptional shutdown phases and chromatin remodeling events suggests that chromatin reorganization may play coordinated roles in the progressive establishment of MSCI.

Our analysis of *Topbp1^B5/B^*^5^ spermatocytes supports this stepwise model of MSCI. TOPBP1 is required for ATR activation^45^. In these mutant spermatocytes, the XY body exhibits immature condensation and mispositioning, and these defects coincide with disruption of the second silencing phase, as indicated by persistently high transcriptional activity in late pachytene compared to wild type (Fig. 2C). This suggests that proper maturation of the XY body is required for MSCI completion and highlights the role of sex-body architecture in stabilizing transcriptional silencing.

Another important contribution of our study is the characterization of the biophysical properties of the mature sex body. Previous studies have proposed that phase separation may underlie the formation of the XY body, enabling silencing through the establishment of a dense, membrane-less compartments^29,30^. While these hypotheses were largely conceptual, our data provide supporting evidence through live-cell dye exclusion assays. We found that mature sex bodies resisted dissolution by LLPS-disrupting chemicals, indicating that they adopt a reversible gel-like state, contrasting with the fully liquid-like phase-separated compartments proposed by Lin et al. (2024) and Xu and Qiao (2021)^30,34^. This condensed state aligns with a “molecular exclusion” model, where limited permeability may prevent entry of small molecules and, by extension, larger regulatory factors such as RNA Pol II. Thus, it is reasonable to propose that gel-like phase separation promotes robust transcriptional silencing in mature sex bodies by establishing a restrictive physical barrier. However, whether this phase transition is sufficient to enforce MSCI remains to be demonstrated. Future studies should specifically test whether disrupting phase separation alters both molecular permeability and transcriptional repression, thereby establishing a causal link between these processes.

Beyond MSCI, we extended our analysis to the broader context of meiotic silencing, particularly MSUC. Although MSUC has been reported to transcriptionally silence unsynapsed chromatin, its extent has not been precisely quantified either cytologically or transcriptionally. To address this gap, we generated a *Spo11^−/−^* scRNA-seq library to serve as a well-established MSUC-positive model. Our results revealed a small subpopulation of “pachytene-checkpoint-escaping” cells that express high levels of post-meiotic (spermatid-stage) markers such as *Tnp1/2*, *Prm1/2* (Figs. 4B,C). Although these 75 cells represent a minor fraction of the *Spo11^−/−^* spermatocytes, we observed “mature” pseudo sex bodies within this population (Fig. 4E). Notably, the field remains divided on whether MSUC universally results in silencing. Some studies have suggested that MSUC can occur in γH2AX-labeled pseudo sex bodies^36^, while others argue that unsynapsed X chromosomes are not silenced, such as in *C. elegans*^46^. Here, our findings provide direct evidence for transcriptional silencing within these *Spo11^−/−^*SPT-like spermatocytes, supporting the presence of MSUC in mammalian spermatocytes. Biologically, this underscores the complexity and potential evolutionary divergence of MSUC regulation, adding to the ongoing discussion that the extent and mechanisms of MSUC may vary by species or cellular contexts.

Additionally, our use of three distinct synapsis-defective models provides valuable insights into how the extent of synaptic failure shapes MSUC outcomes. In *Spo11^−/−^* mutants, which lack programmed DSBs and show widespread asynapsis^3^, we observed robust silencing across both sex chromosomes and autosomes. *Tardbp* and *Nelfb* mutants displayed more moderate synapsis defects. These differences suggest that MSUC scales with the severity of asynapsis. Moreover, MSUC is hypothesized to share silencing mechanisms with MSCI, as both processes involve similar upstream chromatin-modification signals, such as BRCA1, ATR, and γH2AX phosphorylation^43,47^. However, whether MSUC also relies on downstream mechanisms such as chromatin condensation and phase separation–hallmarks of MSCI–remains unclear. In our study, we found that pseudo sex bodies can undergo transcriptional silencing despite being structurally distinct from canonical sex bodies. These pseudo sex bodies were often larger, less condensed, and centrally located within the nucleus (Fig. 4E), yet still exhibited MSCI-like repression (Figs. 4H,I). This finding suggests that MSUC does not strictly require chromatin condensation or spatial sequestration to repress transcription, highlighting a mechanistic divergence from MSCI. While previous studies in XO oocytes showed that γH2AX-positive domains gradually condense over time and eventually exclude Cot-1 signals during pachytene^13^, this likely reflects the behavior of a single unsynapsed sex chromosome and is more analogous to entire X chromosome silencing in MSCI than to the partial chromosome silencing of MSUC. Together, these observations raise the possibility that MSUC may employ alternative or additional silencing strategies beyond those used in MSCI, opening new avenues for investigating the stepwise and context-dependent mechanisms that govern transcriptional silencing across different synapsis and chromosomal landscapes.

Finally, it has been hypothesized that MSUC in oocytes, analogous to MSCI in spermatocytes, might serve as a meiotic checkpoint to eliminate defective oocytes and thereby contribute to oocyte quality control. Silencing essential genes for oogenesis via MSUC could induce apoptosis, potentially explaining the pronounced oocyte attrition around birth. Using our digital chromosome banding method, we were able, for the first time, to precisely evaluate MSUC’s potential contribution to oocyte attrition. Unexpectedly, analysis of 456 late-pachytene oocytes revealed no MSCI-like severe silencing, suggesting a sexual dimorphism in asynapsis-induced meiotic silencing.

## Methods

### Mice

All mice used in this study were from a C57BL/6J background. They were housed in the College of Veterinary Medicine Animal Care Facility at the University of Illinois Urbana-Champaign (UIUC), and maintained in accordance with the guidelines established by the UIUC Institutional Animal Care and Use Committee (IACUC). The *Spo11* mutant line and its corresponding genotyping procedure has been previously described in detail^2^. SCML2-mClover mice were generated by fusing the *mClover* sequence to *Scml2*, enabling visualization and tracking of SCML2 protein expression and localization^31^.

### Immunofluorescence microscopy

The surface spreading of spermatocyte nuclei and immunostaining procedure were performed as previously described^48^. For immunostaining, the following primary antibodies were incubated overnight at room temperature: rabbit anti-SYCP3 (ab15093 Abcam, 1:200 dilution), mouse monoclonal anti-γH2AX (05-636 Millipore, 1:500 dilution). Subsequently, the slides were incubated for 1 hour at 37°C with the corresponding goat secondary antibodies: Alexa Fluor™ 488-conjugated goat anti-mouse IgG (H+L) (Fisher Scientific, 1:1000), Alexa Fluor™ 594-conjugated goat anti-mouse IgG (H+L) (Fisher Scientific, 1:1000), Alexa Fluor™ 488-conjugatd goat anti-rabbit IgG (H+L) (Fisher Scientific, 1:1000), Alexa Fluor™ 594-conjugated goat anti-rabbit IgG (H+L) (Fisher Scientific, 1:1000). Coverslips were mounted with Prolong Gold antifade reagent premixed with DAPI (1µg/ml). All of the immunofluorescence images (Including the Z-stack) were captured using a Nikon A1R confocal microscope and processed by NIS-Elements software.

### Nuclear positioning measurement of sex bodies

Stage-specific images of wild-type spermatocytes were examined to distinguish immature from mature sex-bodies. Different image channels were merged and analyzed using ImageJ (version 1.53v). ImageJ’s enhancement features, including ‘smoothing’, ‘noise reduction’, and ‘edge finding’, were applied to improve image clarity. To compare the localization of sex bodies, we employed ImageJ’s ‘freehand line tool’ to measure two distances: ‘R’ and ‘r’ (Extended Data Fig. 6). The distance ‘r’ represents the distance between the center of the cell nucleus and the center of the sex body, whereas ‘R’ extends from the center to the outer-edge of the nucleus. A ratio of ‘r’ over ‘R’ provided an indication of how close the sex body was to the center of nucleus.

### 1,6-hexanediol and ThT treatments

To test whether matured sex bodies exhibit liquid-like structures, as indicated by Lin et al., (2024)^34^, SCML2-mClover spermatocytes were stained with 0.1 µg/ml Hoechst 33342 (Sigma-Aldrich) to visualize DNA in living cells. The same spermatocyte was imaged before and after a 20-minute treatment with 5% 1,6-hexanediol, a compound that disrupts weak hydrophobic interactions in liquid-like structures. This allowed us to directly examine the effect of hexanediol on sex bodies within the same nucleus (Fig. 3D).

To test whether mature sex bodies form amyloid solid-like structures, we used Thioflavin T (ThT, Fisher Scientific, NC0517036), a small molecular dye which emits green fluorescence and is commonly used to stain DNA and irreversible protein aggregates. First, we visualized the relative position of mature sex bodies using SCML2-mClover green fluorescence within the Hoechst-stained nuclei of live spermatocytes. To avoid interference from the overlapping green fluorescence of ThT and mClover, the SCML2-mClover signal was quenched by exposing the sample to a 488 nm laser for 5 minutes using a Nikon confocal microscope. After quenching, the same late pachytene/early diplotene spermatocyte nuclei were stained with 0.05% ThT and immediately imaged. This approach allowed us to examine whether mature sex bodies adopt a solid-like structure (Extended Data Fig. 10).

### Evaluation of DNA dye (Hoechst 33342 and DRAQ5) penetration in spermatocyte nuclei

Based on the dynamic localization of SCML2 during prophase I, SCML2 nuclear patterns can help stage and mark sex bodies. SCML2 is widely distributed throughout the nucleus before pachynema, but later becomes concentrated in the sex-body region during pachynema and diplonema. This transition in SCML2 localization serves as a useful marker for assessing sex-body maturation^31^. A fully concentrated SCML2 signal indicates a mature sex body, whereas a more dispersed SCML2 signal suggests an early prophase I stage, potentially containing an immature sex body.

To assess the penetration velocity of DNA dyes in spermatocyte nuclei, Hoechst 33342 (0.1 µg/ml) was added to SCML2-mClover spermatocyte suspensions, and images were captured at various time intervals (5 min, 15 min, 25 min). At 5 minutes, weak signals in the nuclear interior indicated that Hoechst had not yet fully penetrated the nuclear center (Extended Data Fig. 9). By 15 minutes, Hoechst intensity increased in the sex body and inner nuclear regions, while staining in the outer nucleus (excluding the sex body) remained relatively unchanged (Fig. 3A and Extended Data Fig. 9).

To quantify Hoechst penetration into the sex body, we calculated the relative Hoechst staining by dividing the signal intensity of the sex-body region by that of the outer nuclear region (excluding the sex body). Staining values were determined at 5, 15, and 25 minutes after Hoechst addition (n=40, 33, and 42, respectively) (Fig. 3B). The relative Hoechst staining increased significantly after 25 minutes, indicating that Hoechst does not rapidly enter the sex-body regions in live spermatocytes. To confirm this result, we repeated the experiment using another small DNA dye, DRAQ5 (Novus Bio) (in 1:500) (Fig. 3C).

### scRNA-seq library construction

Wild-type and *Spo11^−/−^* mutant testes were dissected and placed in ice-cold PBS buffer. The tunica albuginea was carefully removed, and the seminiferous tubules were transferred to digestion buffer 1 (2mg/ml Collagenase I, 10% KSR in α-MEM). Subsequently, the tissue was shaken to facilitate cell release. The sample was then incubated in a 34°C water bath while rotating at 215 rpm for 30 min. Every 5 min, a wide pore pipette was used to gently pipette the sample up and down five times to further dissociate cells. To enhance single-cell yield, the sample was washed with 1X HBSS and then immersed in digestion buffer 2 (700 µg/ml trypsin dissolved in HBSS) for further digestion at 34°C and 215 rpm for another 10 minutes. The resulting cell suspension was filtered through a 40-µm cell strainer, washed with HBSS, and resuspended in 1ml of HBSS containing 0.04% BSA. The scRNA-seq library construction was performed in the Roy J. Carver Biotechnology Center at UIUC using 10X Genomics platform. Next, the library was sequenced on a single lane of the NovaSeq SP system using 2 × 150 nt sequencing.

### scRNA-seq data analysis

WT and *Spo11^−/−^* raw sequencing data were mapped to the mm10 reference genome using CellRanger (4.0.0). The resulting gene count matrices and barcode information were further analyzed in R (4.2.1) using the Seurat package (4.3.0). To ensure high data quality, we first removed low-quality cells with a high percentage of mitochondria genes or a low number of expressed genes. After filtering, cells were normalized, scaled, and clustered using the default setting of Seurat. In wild-type mice, a total of 1,426 germ cells were obtained. Based on cell-type-specific markers, these cells were classified into spermatocytes, spermatids, and spermatozoa (Extended Data Figs. 1A,B). In comparison, 4,283 testicular cells were collected from *Spo11^−/−^* mice. Among them, 274 spermatocytes were identified and collected for further analysis (Figs. 4A,B; Extended Data Figs. 11 and 12A).

To refine substage identification, germ cells were subsetted, re-normalized, re-scaled, and re-clustered using a panel of stage-specific markers, including *Stra8*, *Dmrt1*, *Ccnd1*, *Plk1*, *Esx1*, *Prss50*, *Smc3*, *Sycp3*, *Sycp1*, *Crabp1*, *Meiob*, *Dazl*, *Hormad1*, *Hormad2*, *Meioc*, *Piwil2*, *Clgn*, *Prm1*, *Tnp1,* and *Tnp2.* Clustered cells were visualized by UMAP (Extended Data Fig.1). For previously published scRNA-seq datasets (*Topbp1^B5/B^*^5^ and *Topbp1^+/+^*, *Nelfb* cKO, and *Tardbp* cKO, see Supplementary Table 1), we followed similar analytical procedures as described for *Spo11^−/−^*. For oocyte scRNA-seq dataset analysis (see Supplementary Table 1), additional steps were taken. Doublets were identified and removed by using the DoubletFinder (2.0.3). Batch effects were removed and multiple datasets were integrated by applying the Harmony algorithm. Oocyte substages were identified using a set of gene markers, including *Utf1*, *Pou5f1*, *Stra8*, *Rec8*, *Rad51ap2*, *Mei4*, *Msh5*, *Ugt8a*, *Meioc*, *Dmc1*, *Spdya*, *Spo11*, *Ly6k*, *Meiob*, *Mlh1*, *Mlh3*, *Msh4*, *Topaz1*, *Ybx2*, *Sycp2*, *Gtsf1*, *Sycp3*, *Sycp1*, *Syce3*, *Taf7l* (Extended Data Fig. 3B).

To minimize the influence of RNA stability and better estimate newly synthesized transcripts, we analyzed intronic reads representing nascent pre-mRNA using velocyto-generated loom files (loomR v0.2.1.9000). Unspliced reads were extracted and matched to cells in the Seurat object based on shared barcodes. For each cell, total pre-mRNA abundance was calculated as the sum of intronic reads across all genes. Intronic reads from X-linked genes were further quantified to evaluate X chromosome transcription across spermatocyte substages.

### snRNA-seq data analysis

A previously published mouse testis single-nucleus RNA-seq dataset^19^ was obtained from the public repository. Raw count matrices were processed in R (v4.2.1) using the Seurat package (v4.3.0). Data were normalized, highly variable genes were identified, and the data were scaled followed by principal component analysis (PCA). Cells were clustered using the shared nearest neighbor algorithm and visualized by UMAP. Germ cells were identified based on the expression of germline markers including Ddx4, Dazl, Sycp1, Sycp3, and Piwil1, and were subsetted for downstream analysis. Developmental trajectories were inferred using the Slingshot (v2.6.0) algorithm based on PCA embeddings. Genes located on the X chromosome were defined using a curated mouse X-linked gene list, and the log10 ratio of X-chromosome to autosomal expression (X/A) was calculated for each cell along the pseudotime-ordered germ cell trajectory.

### Digital-chromosome-banding analysis

In the digital-chromosome-banding analysis, each chromosome was partitioned into consecutive 10-Mb bins from the 5’ to the 3’ end, with genes allocated to these bins according to their genomic locations. For the scRNA-seq data, transcription counts for each gene were first log10-transformed and then scaled by a factor of 10,000 to reduce variation across cells. The scaled counts from genes within the same bin were summed up to compute the total transcription for each bin, which reflects the bin-level transcription. The matrix was further log-transformed for visualization in heatmaps using pheatmap package (3.17).

## Data availability

The single cell RNA sequencing data (WT and *Spo11^−/−^*) generated in this study have been deposited in the NCBI GEO database under accession number GSE233871. All publicly available sequencing datasets analyzed in this study are listed in Supplementary Table 1.

## Code availability

The source code for scRNA-seq data analysis, MSUC analysis, figure generation are publicly available via GitHub at https://github.com/Qiaolab1/digital-chromosome-banding.

## Author Contributions

YP, SW, RS, XT, and EZ conducted the experiments. SW, YP, XT, EZ, and RS conducted the data analysis. SW, YP, SN, and HQ contributed to writing and editing of this article.

## Acknowledgement

This work was supported by NIH R00 HD082375, NIH R01 GM135549, and Cancer Center at Illinois.

