## Supplemental Figures for "Digital Chromosome Banding Reveals Distinct Spatiotemporal Dynamics and Sexual Dimorphism in Meiotic Silencing"

**Supplementary Tables**

Supplementary Table 1. Single-Cell RNA-seq data used in this study.

| Dataset ID | SRR accession | GEO accession | Strain | Tissue | Developmental stage | Genotype |
| --- | --- | --- | --- | --- | --- | --- |
| 1 | This study | GSE233871 | C57BL/6 | Testis | Postnatal 6-8 weeks | WT |
| 2 | This study | GSE233871 | C57BL/6 | Testis | Postnatal 6-8 weeks | *Spo11^-/-^* |
| 3 | SRR24572539  SRR24572541 | GSE232588 | C57BL/6 | Testis | Postnatal 8-12 weeks | *Topbp1^B5/B5^* |
| 4 | SRR24572540 SRR24572542 | GSE232588 | C57BL/6 | Testis | Postnatal 8-12 weeks | *Topbp1^+/+^* |
| 5 | SRR24042820  SRR24042821 | GSE228454 | C57BL/6 | Testis | Postnatal day 24 | *Nelfb* cKO |
| 6 | SRR24042818  SRR24042817  SRR24042816 | GSE228454 | C57BL/6 | Testis | Postnatal day 24 | *Tardbp* cKO |
| 7 | SRR8753680 | GSE128553 | C57BL/6 | Ovary | Embryo day E13.5 | WT |
| 8 | SRR8753681 | GSE128553 | C57BL/6 | Ovary | Embryo day E14.5 | WT |
| 9 | SRR8946224 | GSE130212 | C57BL/6 | Ovary | Embryo day E16.5 | WT |
| 10 | SRR12102136 | GSE136441 | C57BL/6J | Ovary | Embryo day E18.5 | WT |

**Extended Data Figs.**

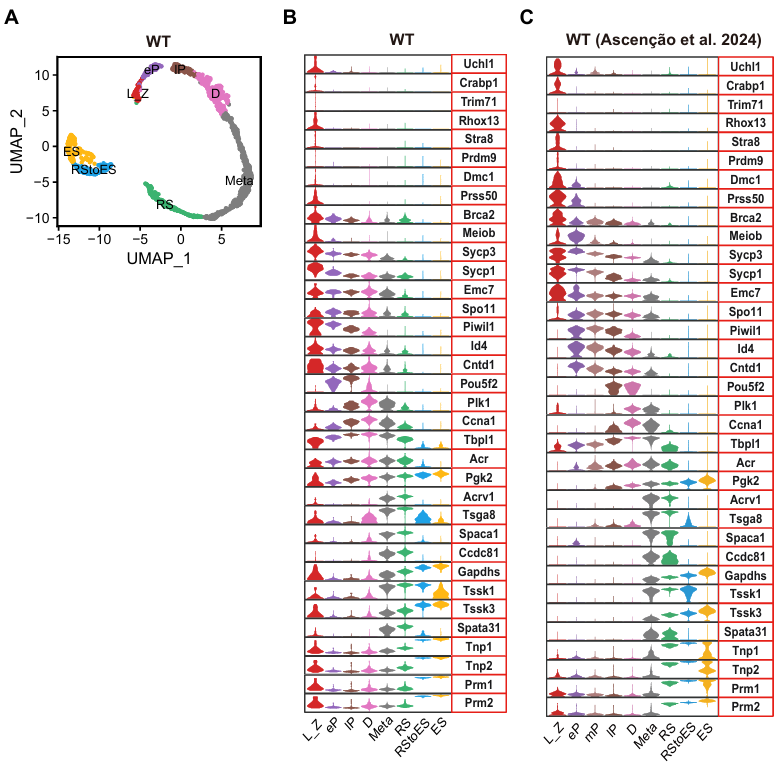

**Extended Data Fig. 1. Stage identification of germ cells in wild type male mice.**

(A) UMAP analysis of 1426 germ cells collected from WT male mice across different developmental stages.
(B-C) Violin plots showing the expression of stage-specific markers across each germ cell stage in wild-type testis, as presented in this study (B) and in Ascenção et al. (2024) (C). L: leptotene; Z: zygotene; eP: early pachytene; mP: mid pachytene; lP: late pachytene; D: diplotene; Meta: metaphase; RS: round spermatids; ES: elongated spermatids.

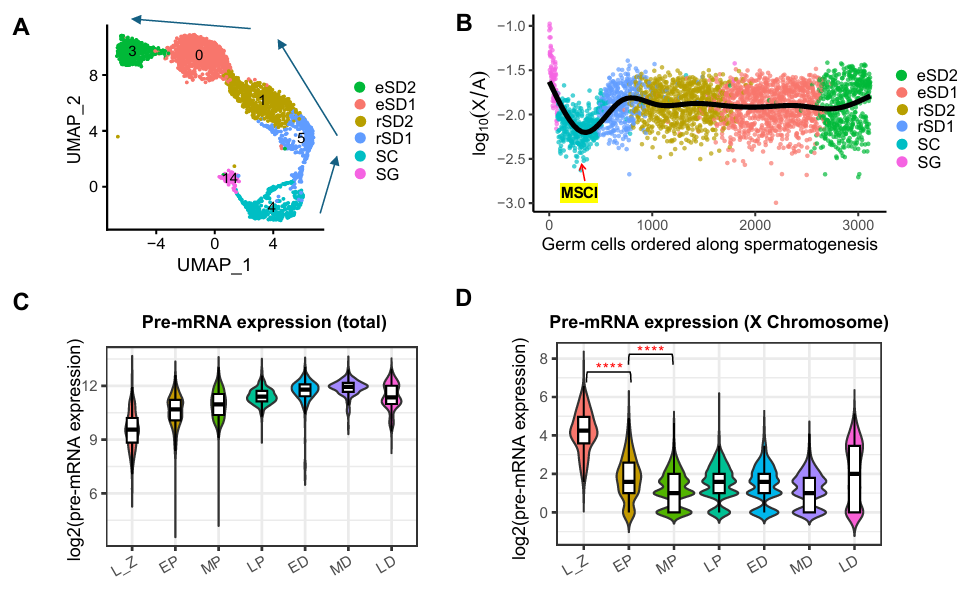

**Extended Data Fig. 2. snRNA-seq and intronic-read analyses of scRNA-seq support progressive X-chromosome transcriptional silencing during meiotic prophase I.**

(A) UMAP visualization of WT mouse germ cells from snRNA-seq dataset, ordered along spermatogenic progression. Arrows indicate the inferred developmental trajectory from spermatogonia to spermatocytes and spermatids. Cell states are annotated as SG, spermatogonia; SC, spermatocytes; rSD1 and rSD2, round spermatids; eSD1 and eSD2, elongating spermatids.
(B) X-to-autosome transcript ratios in individual germ cells ordered along spermatogenic progression. Red arrow indicates MSCI.
(C) Violin plots showing total pre-mRNA expression levels across spermatogenic stages, estimated from intronic reads. Total pre-mRNA levels increase during meiotic progression.
(D) Violin plots showing X-linked pre-mRNA expression levels across spermatogenic stages, estimated from intronic reads. P-value were calculated using Wilcoxon test. **** indicate p-value < 0.0001.

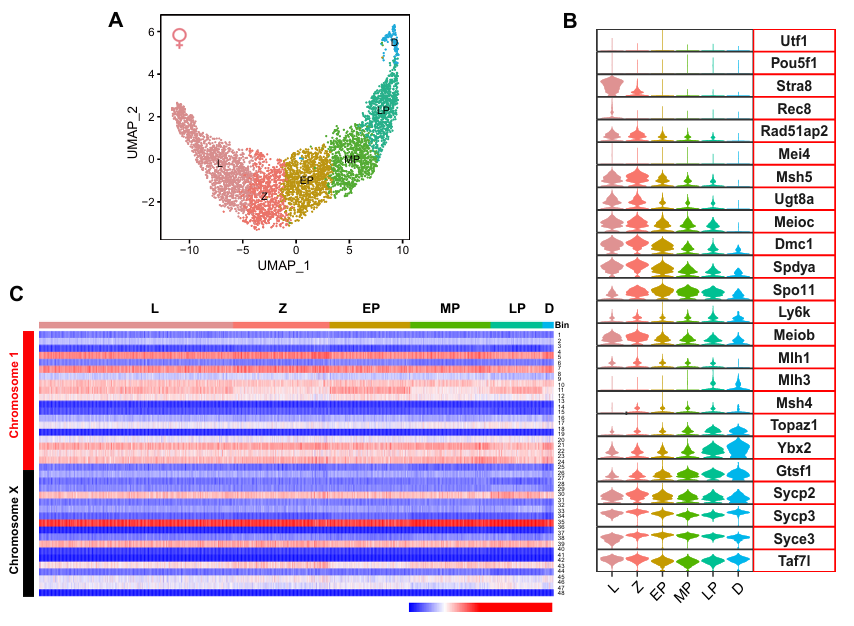

**Extended Data Fig. 3. Digital chromosome banding analysis of oocytes.**

(A) UMAP analysis of 5107 oocytes during meiotic prophase I. We clustered 2078 oocytes for leptonema, 841 oocytes for zygonema, 798 oocytes for early pachynema, 797 oocytes for mid-pachynema, 456 oocytes for late pachytene, and 137 oocytes for diplonema.
(B) Violin plots showing the expression of stage-specific markers across ovarian meiotic prophase I.
(C) Digital-chromosome-banding heatmap of chromosomes X, and 1 across prophase I. Generated from scRNA-seq data. Rows represent bins, while columns correspond to individual cells.

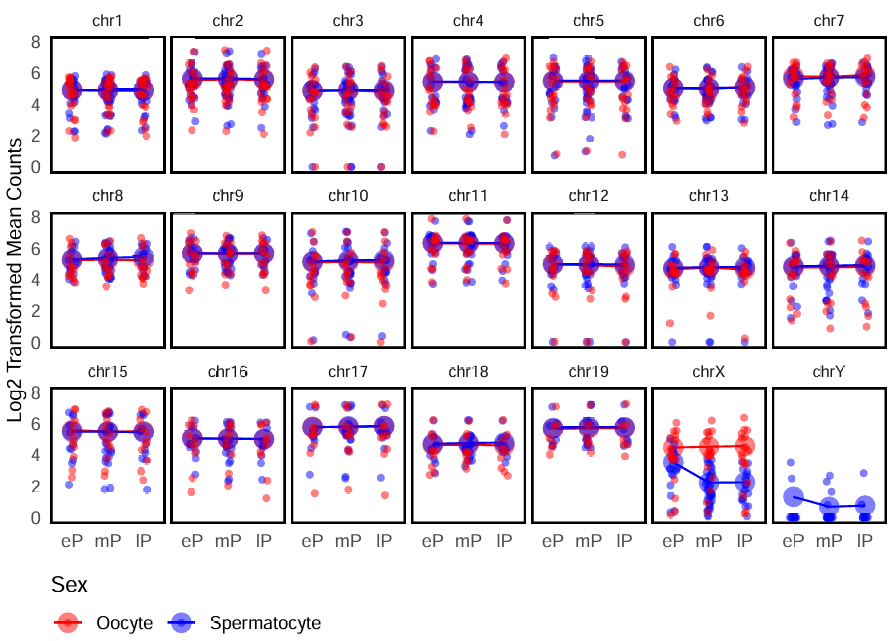

**Extended Data Fig. 4. MSCI is restricted to sex chromosomes in male.**

Normalized transcript counts of each bin (10Mb) on each chromosome in WT pachytene oocytes (in red) and spermatocytes (in blue). Y axis represents the normalized and log-transformed average transcript counts of each bin. Each dot corresponds to one bin along each individual chromosome. X axis stands for three pachytene substages (eP, mP, lP). eP: early pachytene; mP: mid pachytene; lP: late pachytene. Notably, the bins along the X chromosomes of oocytes did not exhibit significant transcriptional repression throughout pachytene substages compared to what was observed in spermatocytes. Spermatocyte X and Y chromosomes undergo MSCI.

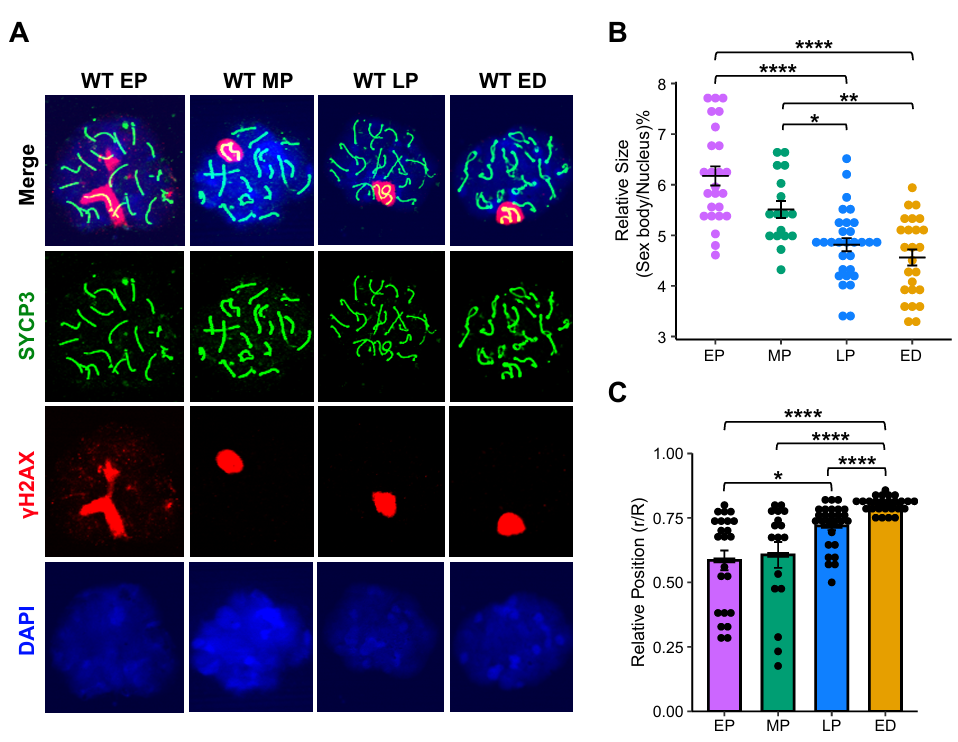

**Extended Data Fig. 5. Wide-type sex bodies maturation in size and nuclear position.**

(A) Immunostaining of chromosome spreads from WT spermatocytes at different substages of prophase I, including early pachynema (WT EP, n=25), mid pachynema (WT MP, n=17), late pachynema (WT LP, n=30) and early diplonema (WT ED, n=25). The spreads were immunostained with γH2AX (red) and SYCP3 (green). The nuclei were counterstained with DAPI. Scale bar = 25 μm.
(B) Comparison of the relative size among sex bodies at different stage. The Y axis represents the relative size, calculated as the area ratio of the sex body to the nucleus. P-value were calculated using t-test. *, ** and **** indicate p-value < 0.05, 0.01 and 0.0001, respectively.

(C) Comparison of the nuclear positioning among sex bodies at different stage. Distances from the center of sex bodies to the center of nuclei were measured. Mature sex bodies in late pachynema are significantly farther from the nucleus center compared to immature sex bodies in early pachynema. Distance ‘r’ represents the distance from the center of the nucleus to the center of the sex body, whereas ‘R’ stands for the nucleus radius (Extended Data Fig. 5). P-value were calculated using Wilcoxon test. *, and **** indicate p-value < 0.05 and 0.0001, respectively.

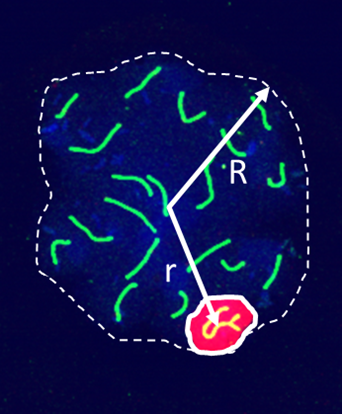

r/R

**Extended Data Fig. 6. Measurement of nuclear positioning of sex bodies.**

A schematic illustrating the measurement of sex body positioning in the nucleus. The distance ‘r’ was measured from the center of the cell to the center of the sex body. and the distance ‘R’ from the center to the edge of the nucleus. The ratio r/R was used to evaluate the localization of sex bodies within the nucleus.

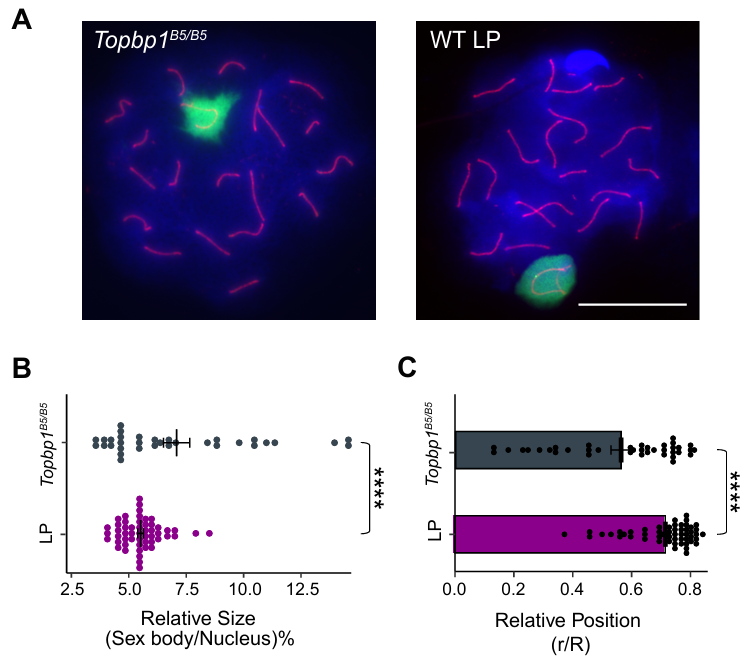

**Extended Data Fig. 7. Comparison of sex-body size and nuclear localization between *Topbp1^B5/B5^* and WT late pachytene spermatocytes.**

(A) Immunostaining of chromosome spreads from *Topbp1^B5/B5^* spermatocytes, and WT late pachytene spermatocytes. Nuclei were stained with SYCP3 (red) and counterstained with γH2AX (green) and DAPI (blue). Scale bar = 25 μm.

(B) Comparison of relative sex-body size in *Topbp1^B5/B5^* spermatocytes and WT late pachytene spermatocytes. P-values were calculated using the Wilcoxon test. **** indicate p-value < 0.0001.

(C) Comparison of relative nuclear positioning of sex bodies in *Topbp1^B5/B5^* spermatocytes and WT late pachytene spermatocytes. P-values were calculated using Wilcoxon test. **** indicate p-value < 0.0001.

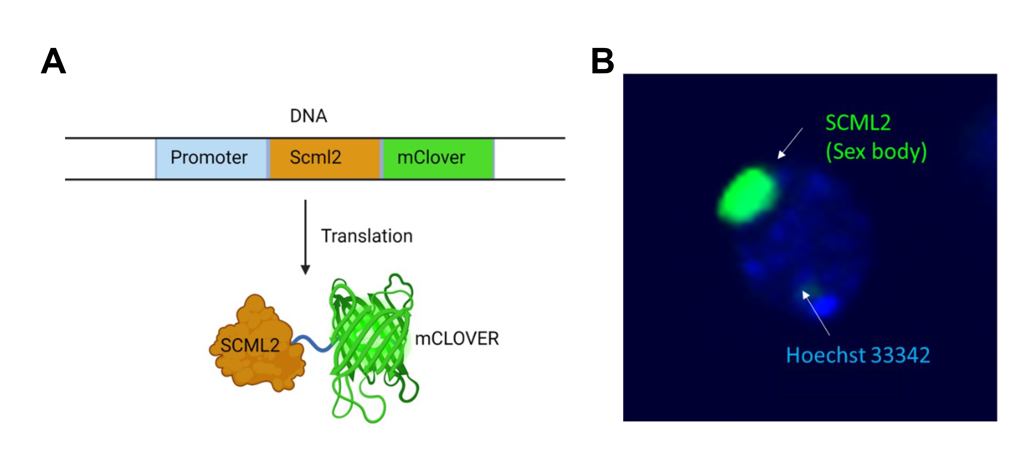

**Extended Data Fig. 8. SCML2-mClover highlights the sex body during prophase I in mouse spermatocytes.**
(A)The SCML2-mClover mouse was generated by adding the mClover DNA sequence to the Scml2 sequence. As a result, the expressed SCML2 is fused with mClover, allowing for the visualization and tracking of SCML2 protein expression and localization.
(B) At pachynema, the SCML2-mClover protein accumulate in the sex body, which highlights the sex body in green.

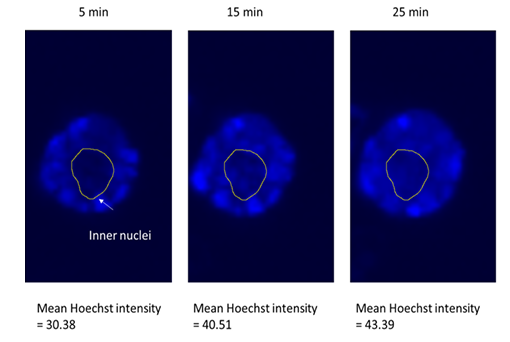

**Extended Data Fig. 9. Hoechst 33342 enter the inner nucleus in living spermatocytes slowly.**

The intensity of Hoechst 33342 staining in the nuclear center of a spermatocyte increased with time, with values of 30.38 at 5 minutes, 40.51 at 15 minutes, and 43.39 at 25 minutes after the addition of Hoechst. The nuclear center is circled by a yellow line in the images.

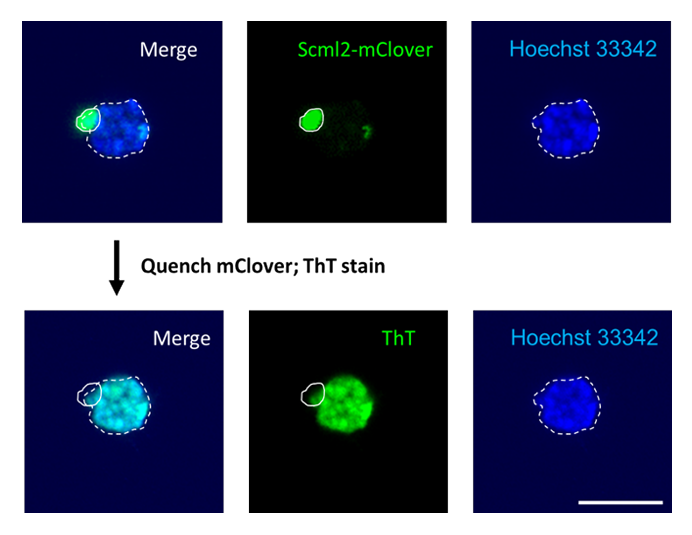

**Extended Data Fig. 10. ThT staining of Scml2-mclover pachytene spermatocyte.**
Hoechst 33342 staining of SCML2-mClover spermatocytes. After imaging, the mClover signal within the same cell was quenched, the cell was stained with 0.05% ThT for taking images as the same position (down). Nuclei stained with Hoechst 33342 was circled by a dash line. The sex body was highlighted by a white line. Scale bar = 25 μm.

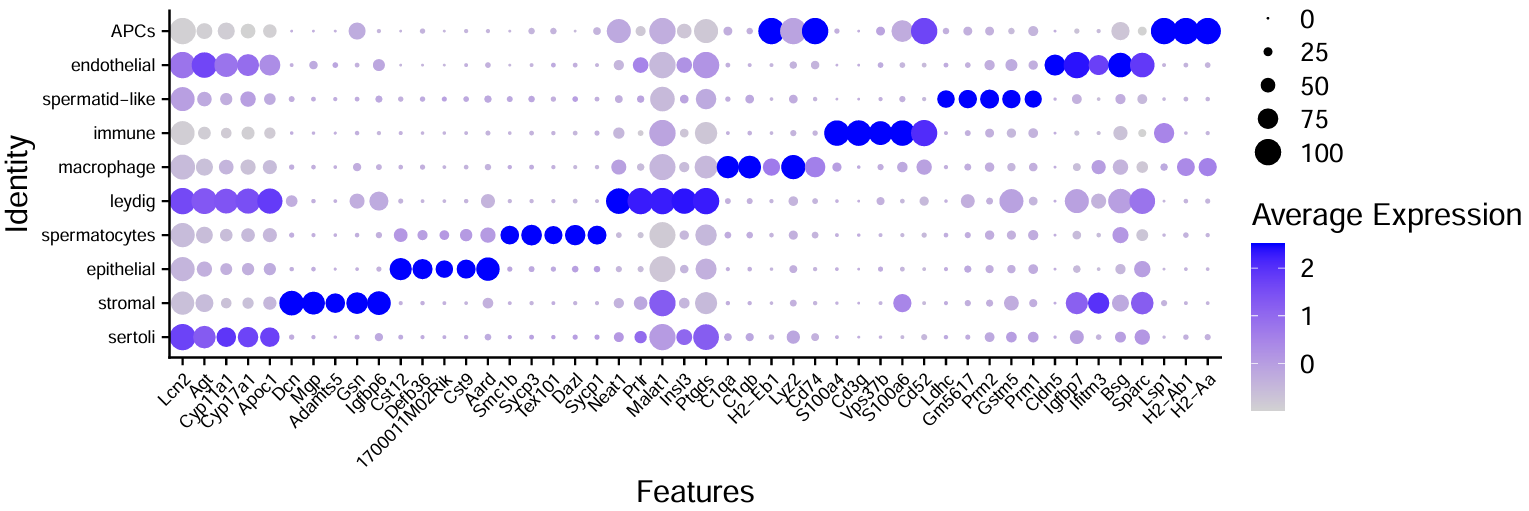

**Extended Data Fig. 11. Cell type identification of *Spo11^-/-^* testicular cells.**
A dot plot showing the average expression levels of selected cell-type-specific markers (column) that were used to identify the clusters for each cell type (row). Dot size indicates the percentage of cells expressing gene marker within the cell cluster.

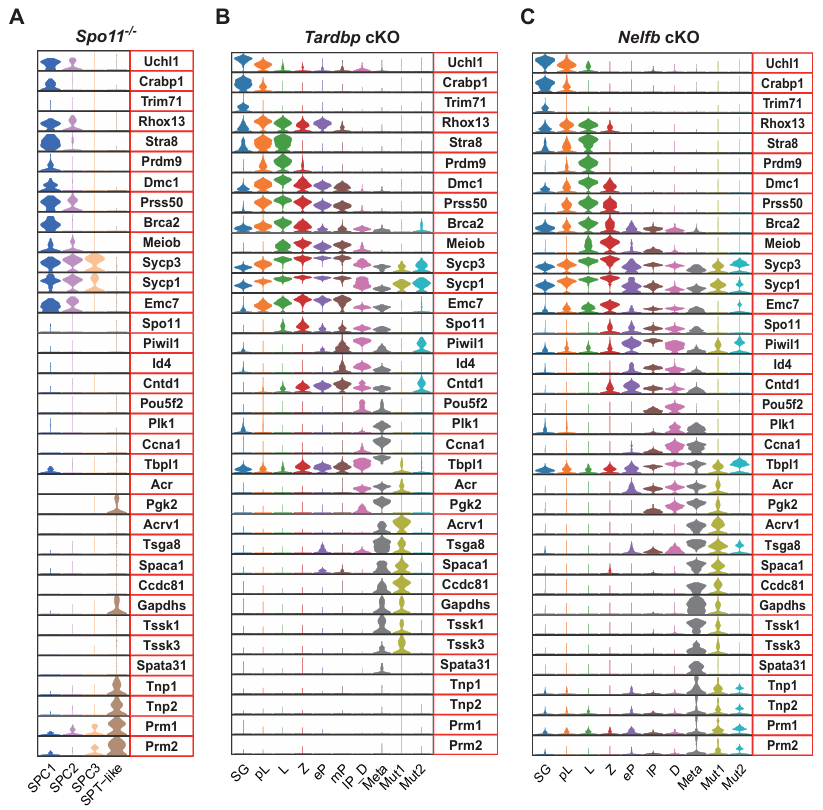

**Extended Data Fig. 12. Stage identification of germ cells in synapsis-defective male mice.**
(A-C) Violin plots showing the expression of stage-specific markers across each germ cell stage in *Spo11^-/-^* (A), *Tardbp* cKO (B), and *Nelfb* cKO (C) testis. SPC1/SPC2/SPC3: spermatocytes; SPT-like: spermatid-like; SG: spermatogonia; pL: pre leptotene; L: leptotene; Z: zygotene; eP: early pachytene; mP: mid pachytene; lP: late pachytene; D: diplotene; Meta: metaphase; Mut1/Mut2: unknown germ cells.

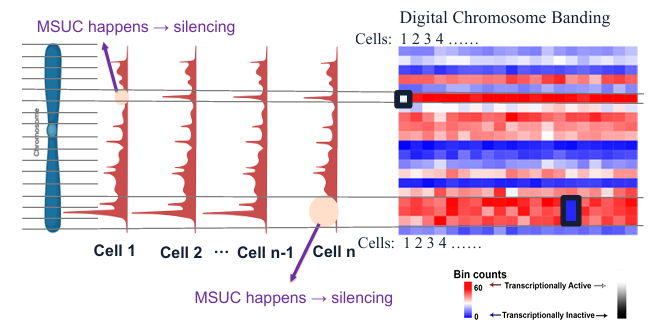

**Extended Data Fig. 13. Schematic of digital chromosome banding for detecting MSUC.**

Each chromosome is divided into 10 MB bins. If MSUC occurs and successfully silences a specific bin, that bin is speculated to show significantly lower transcriptional levels compared to other cell populations.

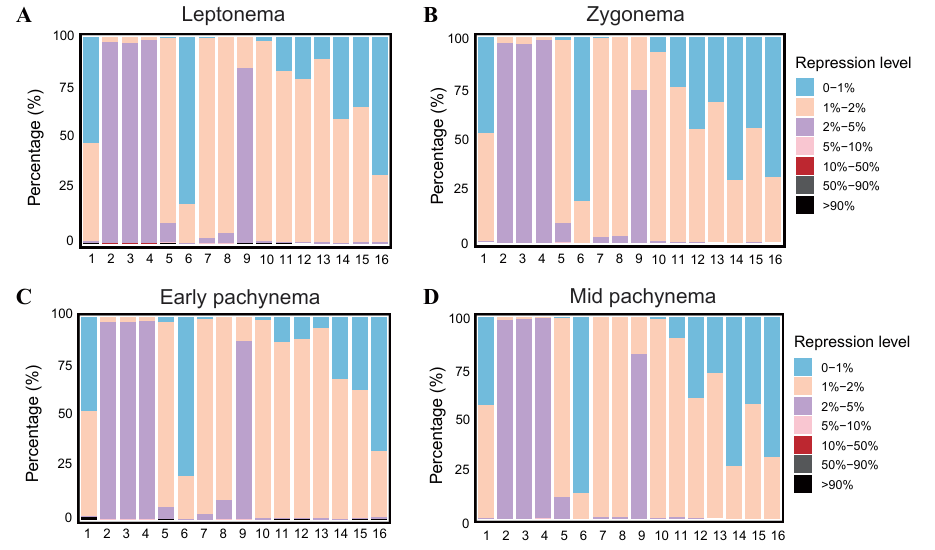

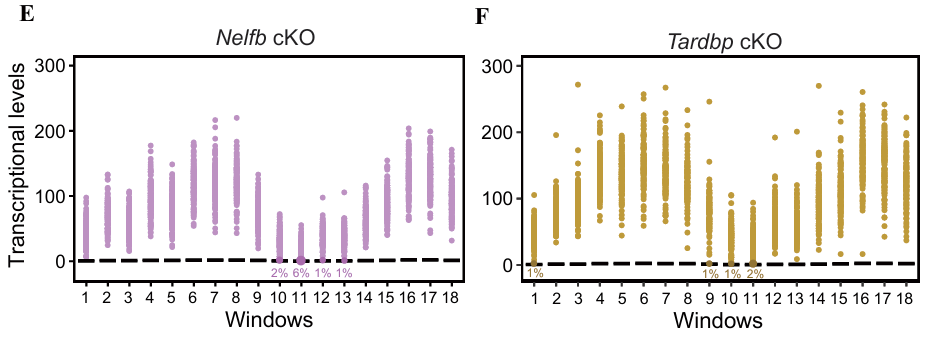

**Extended Data Fig. 14. MSUC exhibits sexual dimorphism between oocytes and synapsis-defective mutant spermatocytes.**

(A-D) Stacked bar plot showing the percentage of transcriptional repression levels in leptotene (A), zygotene (B), early pachytene (C), and mid pachytene (D) oocytes, measured across 16 consecutive 30 MB windows along the X chromosome. Each bar represents a 3-bin (30 MB) window, with colors indicating different levels of transcriptional repression.
(E-F) Graph illustrating transcriptional levels of each 3-bin window along chromosome 1 in *Nelfb* (E) and *Tardbp* cKO (F) mutant spermatocytes. Each dot represents an individual cell. The MSCI full silencing level (100% repression) is defined by the average transcriptional level of WT late pachytene spermatocytes (the stage when full silencing occurs). Black lines indicate MSCI levels for each window. The percentage of cells showing MSCI-like severe silencing is labeled beneath the black MSCI threshold line for each window. Notably, MSCI-like severe silencing is observed in some of the windows.
